# Ants get stuck in traffic jams and resolve them by making adaptive decisions

**DOI:** 10.1101/2023.09.13.557616

**Authors:** Manish K. Pathak, Joy D. Bairagya, Sagar Chakraborty, Sumana Annagiri

## Abstract

Efficient traffic flow underpins biological and engineered systems. Ants, widely regarded as exemplars of collective efficiency in active matter research. We examined ants for traffic dynamics and transport efficiency during nest relocation via tandem running, that affects colony survival. Laboratory experiments, inspired by their monsoon-constrained habitats, and agent-based modeling reveal that narrow paths trigger jams 18 times more frequently than multi-lane paths, primarily driven by returning leaders (59%), followed by brood transporters (27%) and lost followers (14%). Despite a 1.3-fold transport delay, returning leaders mitigate jams through adaptive U-turns and midway recruitment, generating emergent unidirectional flow. This aligns with active matter principles of self-propelled agents navigating non-equilibrium dynamics. These findings reveal traffic jams in tandem-running ants, challenging assumptions of jam-free ant traffic, and inspire algorithms for robotic swarms and traffic optimization. The study contributes to understanding non-reciprocal interactions and emergent phenomena in biological systems.

## Introduction

Traffic flow is critical across diverse systems, from intracellular transport (1) to large-scale animal migrations (2), and even extending to artificial systems like robot swarms and pedestrian traffic(3, 4). Efficient traffic flow is crucial, yet challenging in confined environments, where bottlenecks cause congestion. In human societies, traffic congestion incurs significant socioeconomic costs and low quality of life (5). Similarly, in biological systems, disrupted traffic flow hampers physiological processes and heightens vulnerability to stressors like disease and predation (6–8). These selective pressures that have shaped the evolution of efficient movement strategies across taxa (9). Social insects, particularly ants, are model systems for collective movement. Their self-organized behaviours, driven by simple individual actions, enable efficient collective movement. Ants often exhibit robust traffic flow during foraging, even at high densities, using pheromone trails(10), effective recruitment and lane formation (11). This efficiency has been well-characterized in trail-laying ants (12–15).

However, traffic dynamics beyond foraging remain underexplored. While anecdotal reports describe congestion during cooperative food transport in *Oecophylla smaragdina* (*16*), systematic studies of traffic jams in the other context are scarce. Particularly, bidirectional movement via tandem running— where a leader ant guides a follower through continuous physical contact—is poorly understood (*17*– *20*). Unlike foraging, which involves a subset of foragers, nest relocation requires coordinated movement of the entire colony, including reproductives, non-foragers, callows, and brood. This process faces strong selective pressures, as colony survival depends on its success (*21*–*23*). Environmental perturbations or resource depletion trigger nest relocation, exposing the colony to biotic and abiotic hazards across unfamiliar terrain (*23*). During these relocations, ants may encounter narrow passages, such as those associated with waterlogged areas, dense vegetation and other difficulties on the path that can act as bottlenecks. These constricted paths, in conjunction with the bidirectional flow of ants (i.e., ants in tandem pair moving toward the new nest and returning leaders heading towards the old nest to recruit additional followers), create conditions conducive to traffic jams. Like fire ants excavating narrow tunnels, tandem-running ants during nest relocation face the challenge of navigating confined spaces, making them an ideal system for studying traffic dynamics (*4*). Even though ants are known to have efficient traffic systems, they can have problems in confined spaces. The study of ant traffic dynamics also aligns with the interdisciplinary field of motile active matter, which spans biology, physics, and engineering (*24*). Ants, as self-propelled agents, may exhibit non-reciprocal interactions that drive emergent phenomena.

To address the knowledge gap about traffic management behaviour among tandem running ants in the context of nest relocation in confined space, we combined laboratory experiments on *Diacamma indicum* with agent-based modelling (ABM). The experiments focused on the possibilities of traffic jams at bottlenecks, while the ABM explored the relationship between individual ant behaviours and the emergence of congestion. ABMs provide a valuable tool for studying collective behaviour, enabling systematic manipulation of individual actions and observation of their effects on group-level dynamics (*20, 25*–*32*). Our ABM represented individual ants as agents exhibiting specific behaviours based on experimental data. This allowed us to simulate nest relocations, manipulate individual agent’s behaviours, and quantify their impact on traffic flow. By replicating experimental conditions *in silico*, we aimed to derive mechanistic insights into the processes of potential jam formation and its resolution, that cannot be easily explored from behavioural observations.

## Results

Analysis of 56 jam events in the one-lane path showed returning leaders as the primary cause (59%), followed by brood transporters (27%) and lost followers (14%), a significant deviation from equal distribution (Chi-Square test, *χ*^*2*^ = 17.821, df = 2, *p* < 0.01) highlights the critical role of returning leaders in initiating traffic congestion.

### Path constraints interrupts flow and induces jam

The one-lane path significantly disrupted traffic, causing frequent jams. We observed a substantial increase in tandem run interruptions in the one-lane condition (23.66 ± 10.12) compared to the multi-lane control (8.25 ± 4.71; Wilcoxon paired-sample test, T = 0, n = 12, *p* < 0.01; Fig. 2A), corroborated by the ABM (19.98 ± 7.08 vs. 7.52 ± 6.05; Mann-Whitney U test, *p* < 0.01).

**Fig. 1.**
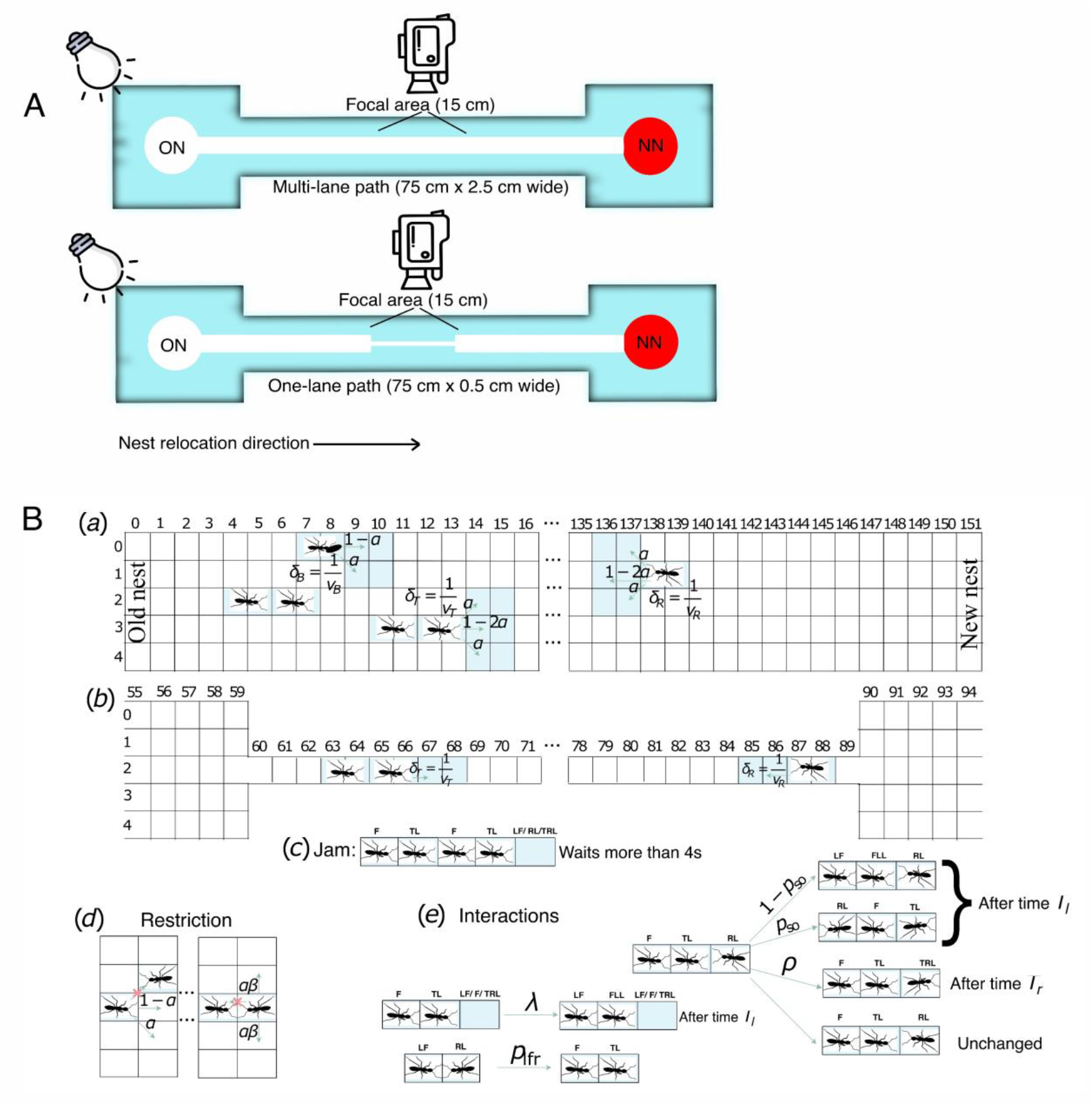
(A) Arena for control and manipulation experiment. (B) Protocol used in the agent-based simulations: (a) and (b) depict the ant’s movement during multi-lane and one-lane experiments. Green arrows show the possible y-directional movements with probability. Along the x-direction, a brood-transporter, a tandem pair and a returning leader move *δB* = 1*/v*_*B*_, *δ*_*T*_ = 1*/v*_*T*_ and *δ*_*R*_ = 1*/v*_*R*_ steps in one second, respectively. Subfigure (c) shows the definition of a jam, the letters F, TL, LF, RL and TRL represent follower, tandem leader, lost a follower, returning leader and turning back returning leader, respectively. Subfigure (d) is the pictorial representation of the fact that two individuals cannot simultaneously occupy the same cell, and the tendency to remain longer in the same position if an ant finds another nest-mate is also introduced in the code by restricting the y-directional movement by a factor *β*. Subfigure (e) depicts the interactions included in the ABM, which are as follows: the creation of a lost follower occurs with a probability of 1 − *p*_so_ during a head-on encounter and with a probability rate of *λ* during a non-head-on encounter; in both scenarios, a minimum time *T*_*l*_ is required to create a lost ant. The switching over happens with a probability of *p*_*so*_ after time *T*_*l*_. The returning leader turns back with a probability rate of *ρ* after time *T*_*r*_, and the recruitment of the lost follower occurs with a probability rate of *p*_*lfr*_ as soon as a returning leader encounters a lost ant.

**Fig. 2.**
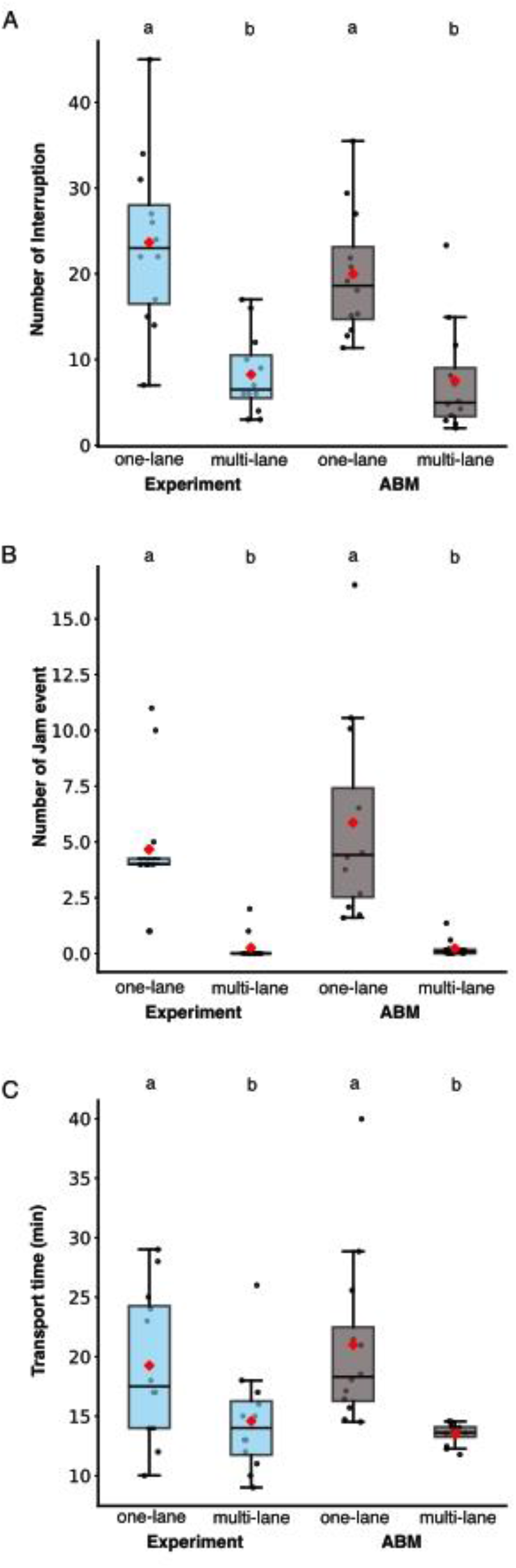
Challenges to tandem running recruitment at a constricted path. **(A)** number of interruptions, **(B)** traffic jams and **(C)** transport time. Comparing multi-lane versus one-lane path in experiment and agent-based model (left and right sections, respectively), experimental observations (blue) and agent-based simulations (grey). Boxes represent interquartile ranges (IQRs), horizontal lines within boxes indicate medians, diamonds represent means, whiskers extend to the 1st and 99th percentiles, and circles show individual data points. Different letters (a vs. b) above the plots indicate significant differences (*p* < 0.05) between one-lane and multi-lane conditions, as assessed by the Wilcoxon paired sample test (experimental data) and Mann-Whitney *U* test (simulation data). We conducted experiments with 12 colonies, each comprising 54–160 ants, and we conducted simulations with 12 colonies of similar size and performed 25 iterations; we used the average for further analysis.

Notably, traffic jams were frequently observed within the one-lane condition, a phenomenon previously unreported in tandem-running ants. Jam frequency was significantly higher in the one-lane setup (4.66 ± 2.99) compared to the multi-lane control (0.25 ± 0.62; Wilcoxon paired-sample test, T = 0, n = 12, *p* < 0.01; Fig. 2B), a result reinforced by the ABM (5.86 ± 4.30 vs. 0.22 ± 0.38; Mann-Whitney U test, *p* < 0.01). Furthermore, both experimental and simulated data revealed a strong positive correlation between colony size and jam frequency (Spearman’s r^2^ = 0.73 and 1.0, respectively; *p* < 0.01), suggesting that larger colonies, with increased ant density, are more susceptible to traffic congestion in confined spaces.

### Impact of jam on transport time

Despite the ants’ ability to navigate through jams, the one-lane path significantly reduced relocation efficiency. The transport phase was prolonged in the one-lane condition (19.25 ± 6.36 min) compared to the multi-lane control (14.58 ± 4.54 min; Wilcoxon paired-sample test, T = 9, *p* = 0.02; Fig. 2C), a result mirrored by the ABM (20.99 ± 7.09 min vs. 13.51 ± 0.86 min; Mann-Whitney U test, *p* < 0.01; Fig. 2C). Furthermore, the frequency of successful tandem runs decreased in the one-lane condition (65.14 ± 12.59% vs. 82.69 ± 9.54%; Wilcoxon paired-sample test, T = 2, *p* < 0.01), consistent with increased interruptions.

Intriguingly, while jam frequency increased 18-fold in the one-lane condition, the overall transport time was only marginally prolonged (1.3 times). This discrepancy suggests that the ants possess a highly effective mechanism for mitigating the detrimental effects of frequent jams, prompting the question: how do they achieve this?

### Jam resolution: Ant’s solution

To dissect the mechanisms by which ants effectively overcome traffic jams, we investigated several potential strategies. Initially, we hypothesized that colonies might reduce the workload by decreasing the number of tandem leaders in the constrained one-lane path. However, the percentage of ants acting as tandem leaders did not significantly differ between the one-lane (20.54 ± 6.69%) and multi-lane (18.07 ± 5.47%) conditions (Wilcoxon signed-rank exact test, V = 16, *p* = 0.07), ruling out this possibility. Similarly, direct observation revealed no evidence of ants stationed to actively regulate traffic flow at the bottleneck, suggesting the absence of dedicated traffic controllers.

Furthermore, we examined whether ants regulated their heading direction in cluster to mitigate congestion. Analysis of ant movement at the bottleneck showed no coordinated flow; instead, approximately equal proportions of ants moved towards the old and new nests (proportion = 0.54), indicating persistent bidirectional traffic. Crucially, the video analysis revealed that jam resolution consistently coincided with ‘mass orientation events’ (Supplementary Video 1), where more than 80% of ants within the jam synchronously aligned towards the new nest, initiating a surge of unidirectional flow. This behavior, observed in 92.85% of jam events, highlighted the pivotal role of mass orientation in resolving congestion.

To elucidate mass orientation, we focused on ‘midway recruitment’ – the recruitment of ants from the path, rather than the nest, by returning leaders. During midway recruitment, a returning leader initiates a new tandem run with a follower ant impeded in the jam or moving undirected after losing its leader, and the pair proceeds to the target nest. We found that midway recruitment of lost followers increased substantially (by 115.38%) in the one-lane relocations (27.53% ± 10.72%) compared to the multi-lane relocations (13.07% ± 7.89%) (Wilcoxon paired-sample test, T = 2, n = 12, *p* < 0.01; Fig. 3A). The simulations mirrored this, showing a similar increase (27.63% ± 2.93%) compared to (9.99% ± 5.13%) (Mann-Whitney U test, n = 12, *p* < 0.01; Fig. 3A). Midway recruitment via switching-over also rose significantly (by 75%) in one-lane relocations (7.31% ± 5.60%) compared to multi-lane (4.23% ± 4.07%) (Wilcoxon paired-sample test, T = 9, n = 12, *p* = 0.02; Fig. 3B), with simulations showing a similar trend (6.68% ± 1.08%) compared to (4.15% ± 1.78%; Mann-Whitney U test, n = 12, *p* < 0.01; Fig. 3B). Overall, returning leaders showed significantly higher involvement in midway recruitment in one-lane relocations (61.19% ± 14.70%) compared to multi-lane (40.38% ± 13.47%) (Wilcoxon paired-sample test, T = 2, n = 12, *p* < 0.01). The finding suggests that midway recruitment at individual level contributes to the emergence of mass orientation at the group level.

**Fig. 3.**
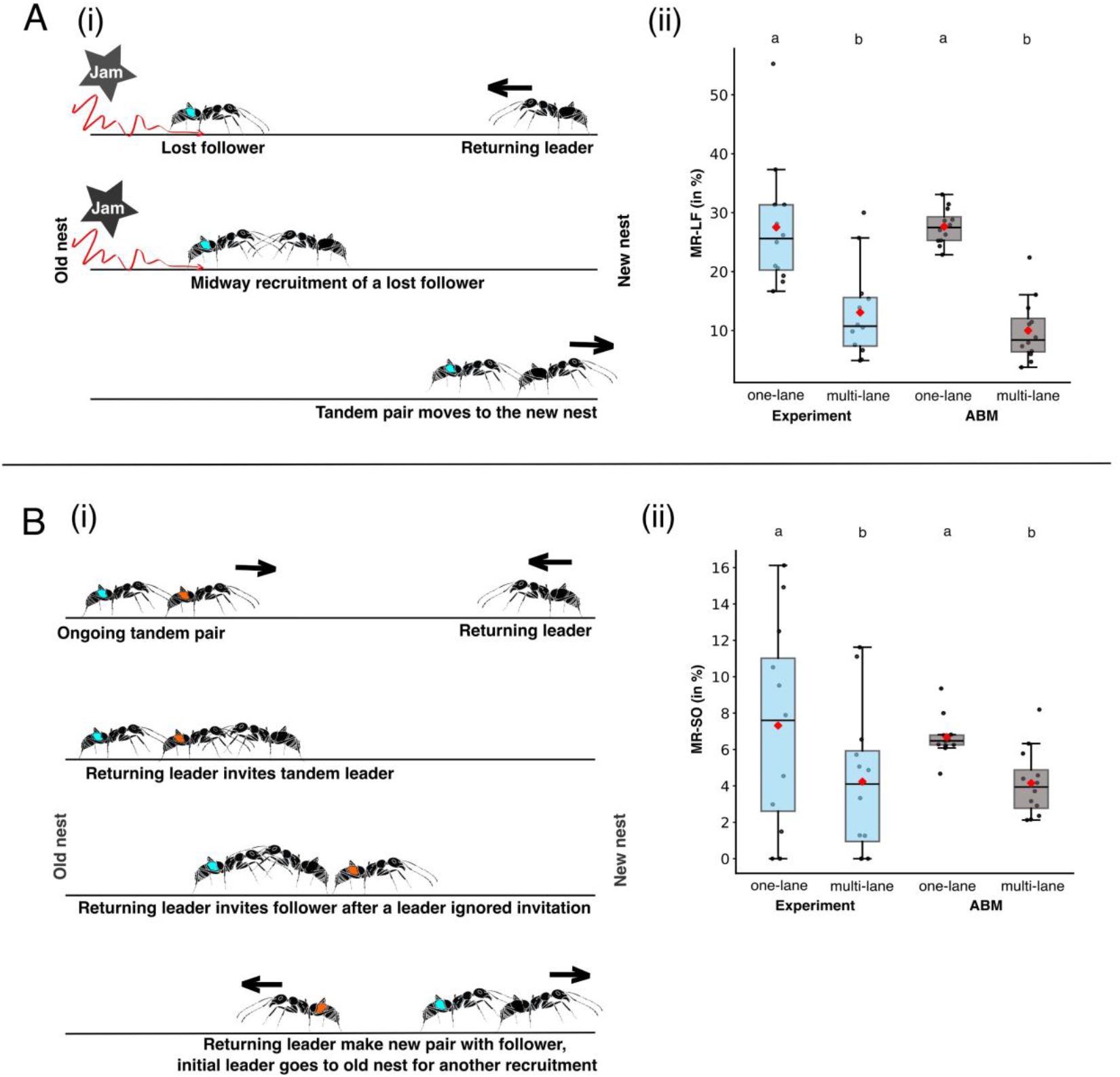
A constricted path increases midway recruitment. **(A) *Recruitment of Lost Followers:*** (i) A schematic illustrates a returning leader recruiting a lost follower from the path. Box-and-whisker plots compare the number of lost follower recruitments (MR-LF) in (ii) one-lane and multi-lane paths based on experimental observations (grey) and simulations (blue). MR-LF events are significantly more frequent in the one-lane path (*P* < 0.05). **(B) *Switching Over:*** (i) A schematic illustrates a returning leader actively breaking an ongoing tandem pair on the path and recruiting the follower (MR-SO). The box-and-whisker plots (ii) of the experimental observations and simulations show a significant increase in midway recruitment by switching over in the one-lane compared to the multi-lane paths (*P* < 0.05).

We used ABM to investigate the causal relationship between midway recruitment (MR-SO and MR-LF) and jam resolution. In simulations, we systematically varied the probabilities of lost follower recruitment (*p*_*lfr*_)and switching-over (*p*_*so*_). MR-LF had a negligible effect on jam resolution (Fig. 4A(ii)), although reducing it increased jam frequency (Fig. 4A(i)). Decreasing (*p*_*lfr*_)also led to an increase in transport time (Fig. 4A(iii)). In contrast, MR-SO positively impacted jam resolution (Fig. 4B(ii)). Increasing (*p*_*so*_) led to faster resolution of more jams which makes transportation faster (Fig. 4B(iii)).

**Fig. 4.**
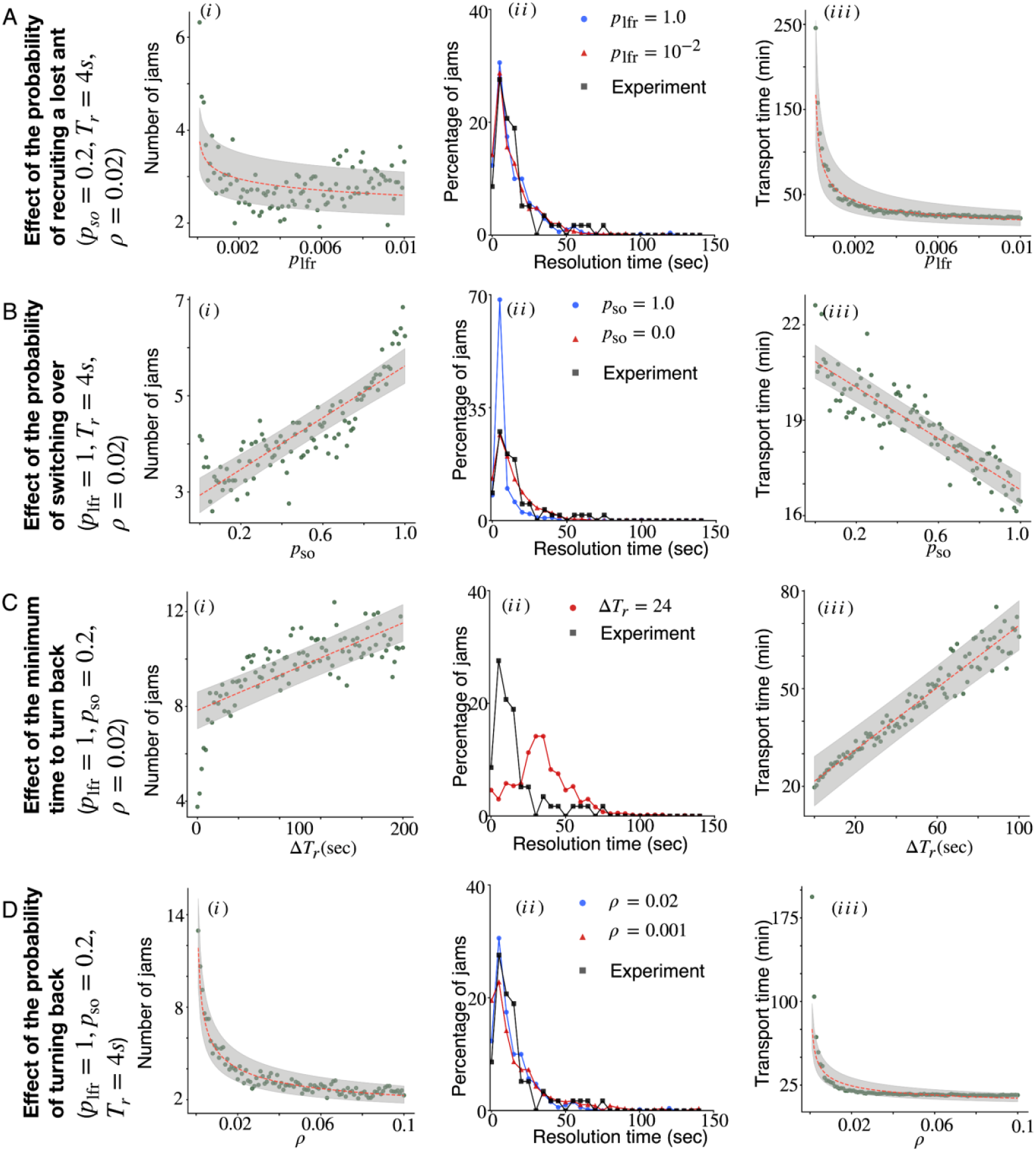
Role of returning leaders’ behavioural flexibilities in jam resolution: Each panel **demonstrates** how (i) the number of jams, (ii) the distribution of jam resolution time, and (iii) the transport time depend on the specific behaviour of the returning leader. Panels **(A), (B), (C)**, and **(D)** show the effect of the probability of recruiting a lost follower (*p*_*lfr*_), the probability of switching over (*p*_*so*_), the minimum time required to turn back (*T*_*r*_), and the probability of turning back (*ρ*) after waiting for (*T*_*r*_) time, respectively. To examine the role of a specific behaviour (e.g., switching over), we fixed all other variables (*p*_*lfr*_, *ρ, T*_*r*_) as mentioned along the *y*-axis while continuously varying the parameter defining that behaviour ((*p*_*so*_), in this case). Additionally, the grey areas in all subplots of the first and third columns represent the 95% confidence intervals of the fitted estimate (red dashed line). These results indicate that the probability of switching over (*p*_*so*_), minimum time required to turn back (*T*_*r*_), and the probability of turning back (*ρ*) are crucial factors in jam resolution, whereas the probability of recruiting a lost follower (*p*_*lfr*_) plays no significant role. Furthermore, all behaviours except switching over influence the number of jams and total transport time. For simulations, we set colony size to the mean size of experimental colonies (87 ants) and averaged results over 25 independent realisations.

We further examined the returning leader’s U-turn behaviour by considering turning time (*T*_*r*_)and turning probability (*ρ*). Higher (*T*_*r*_)and lower (*ρ*)increased both transport time and jam frequency (Fig. 4C, 4D). Importantly, even with constant MR probabilities, reduced (*T*_*r*_)and increased (*ρ*)led to faster resolution. Our ABM demonstrates the crucial role of returning leaders in resolving traffic jams. Their ability to rapidly assess the situation and adaptively turn back by making a U-turn and recruiting nearby nestmates from jam site, is key to mitigating jams and optimizing traffic flow.

## Discussion

Our study reveals that tandem-running ants, *D. indicum*, experience traffic jams during nest relocation, demonstrating that even highly efficient ant traffic systems can be disrupted under challenging environmental constraints (*10, 12*–*14*). This susceptibility mirrors observations in other biological systems characterized by bidirectional movement. Using a combined approach of behavioral observation and agent-based modeling, we demonstrate that ants actively resolve these jams through adaptive decisions made by returning tandem leaders. Upon encountering a jam, these leaders initiate U-turns and recruit stranded nestmates, triggering an emergent mass orientation that results into an unidirectional flow towards the new nest.

The adaptive decisions of returning leaders in *D. indicum*—initiating U-turns and midway recruitment—reflect non-reciprocal interactions within single species system, as commonly seen in the living system due to asymmetry of information transfer (*33*). It can drive emergent phenomena, such as unidirectional flow of individuals. These behaviors align with cognitive, self-steering active particles that adapt motion based on environmental cues (*24*). Our ABM systematically capturing how individual decisions lead to collective outcomes. These findings suggest potential algorithms for micro-robotic swarms, where reinforcement learning could emulate ant-like adaptability to optimize navigation in confined spaces (*24*).

This emergent unidirectional movement, crucial for efficient traffic flow, exemplifies self-organization, where simple individual actions give rise to complex collective outcomes (*1, 4, 13, 16, 34*). This highlights the flexibility of group behaviors in achieving collective goals, consistent with findings in other social systems (*2, 10, 35*). Similar to the ‘backing up’ strategy observed in fire ants navigating confined tunnels (*4*), our research underscores the importance of individual behaviors in regulating collective traffic flow. However, unlike the excavation tasks in fire ants, our study focuses on bidirectional movement during nest relocation, introducing unique coordination challenges.

While feedback and information transfer are pivotal in decentralized systems, including bird flocks and human crowds (*36*–*38*), we observed a distinct lack of other jam-mitigation strategies in *D. indicum*. We found no evidence of workforce reduction, dedicated traffic regulators, or unidirectional cluster-like movement, contrasting with reports in *Lasius niger* (*34*). This suggests that *D. indicum* relies primarily on the adaptive decisions of returning leaders, specifically their U-turn behavior and midway recruitment, to overcome congestion.

Optimal transport is critical during nest relocation, as colony survival and fitness are directly at stake (*23, 37*–*39*). Despite increased jam frequency in the one-lane condition, returning leaders effectively mitigated the impact on overall transport duration. Agent-based simulations further demonstrated the significant influence of returning leaders’ adaptive decisions, such as U-turn timing and switching tendency, on jam resolution time and transport efficiency. This highlights the strong selective pressure favoring strategies that optimize traffic flow and minimize delays during relocation (*42, 43*).

While we did not explicitly construct a fundamental diagram, the observed positive correlation between colony size and jam frequency suggests that *D. indicum* may operate within an optimal occupancy range, akin to findings in other traffic systems.

In conclusion, our study provides novel insights into the self-organized mechanisms by which tandem-running ants resolve traffic jams. We demonstrate that path constraints induce jams, which are subsequently mitigated through emergent mass orientation and midway recruitment, driven by the adaptive decisions of returning leaders. These findings not only address a knowledge gap in the traffic management of tandem-running ants but also offer a compelling model for optimizing flow in diverse systems. The principles observed in *D. indicum* may inspire novel algorithms for applications such as autonomous vehicles and pedestrian evacuation, as well as inform the design of bio-inspired robot swarms (*44, 45*).

Future research should expand on these findings by exploring jam resolution mechanisms in other ant species, particularly those employing chemical trails in non-foraging contexts. Moreover, while our laboratory experiments provided controlled insights, investigations in natural environments are crucial for understanding the full complexity of ant traffic dynamics. Fundamental traffic diagrams can be explored further to elucidate the traffic dynamics in tandem-running ant colonies. One could explore ant-inspired algorithms for micro-robotic swarms (in drug delivery) or pedestrian traffic systems, validating their scalability in active matter systems. Experimental studies should investigate how environmental factors influence ant traffic dynamics, connecting microscopic adaptive behaviors to macroscopic collective outcomes (*24*). These findings advocate for interdisciplinary collaboration to translate these evolutionary insights into real-world applications and advancing the active matter research.

## Materials and Methods

### Model organism and experimental setup

We collected 18 colonies of *Diacamma indicum* ants from their natural habitat in Nadia district, West Bengal, India, using the water flooding method (*46*). Each colony, typically comprising a gamergate (the sole reproductively active female), brood (eggs, larvae, and pupae), and adult female workers, had an average size of 87 individuals (ranging from 54 to 160). Each ant was marked with a unique color for individual identification. Colonies were housed in 9 cm (diameter) Petri dishes modified to serve as artificial nests, red cellophane paper was used for covering the lids to make the nest darker. We made a 1.5 cm entrance hole in each dish and then kept these nests in larger plastic boxes (28.5 x 21.5 x 12 cm) with a plaster of Paris base. Colonies were provided with *ad libitum* water and ant cake (*47*) and allowed them to acclimate in laboratory conditions for 24 hours before experiments began.

We conducted experiments in a rectangular plexiglass arena (100 cm x 40 cm) filled with water (up to 2.5 cm from bottom). Simulating a waterlogged conditions that is commonly faced by these ants during monsoon season in their natural habitat. A new nest box kept 75 cm apart, reflecting typical relocation distances (*18*). A floating plexiglass bridge was placed such that it connects the two nest and serves as the sole path to reach to the other side (Fig. 1A). To find the impact of path width, we used two bridge types: a) muti-lane bridge (control): on which multiple ants can walk in parallel (2.5cm wide); b) one-lane bridge (bottleneck): similar to control bridge except a 15cm long central constriction of 0.5cm width.

### Nest relocation and behavioural observation

Each colony underwent two relocations, control and manipulation in random order, with a 24-hour rest between trials. We cleaned the setup and replaced the water before each relocation. Relocations were initiated by removing the red cellophane from the old nest’s roof and introducing a 9-watt white LED bulb 20 cm above it to induce stress that stimulate few ants for the exploration of habitable shelter in arena. We recorded (a) discovery time (light induced stress to new nest discovery), (b) latency (discovery to first tandem run), and (c) transport time (first tandem run to last ant/brood entering the new nest).

We used a Sony Handycam^®^ (HDR-CX200) video camera to record the ant behaviour during relocations, focusing on a 15 cm long path at the center of bridge and 7.5 cm around it’s edges, where we located the constriction in the one-lane condition (Fig. 1A). We decoded the video-recording for analyzing various aspects of ant’s behaviour, including (a) heading direction (towards the old or new nest) and time of tandem runs and brood transports, (b) interrupted tandem runs, either disrupted or resumed (c) midway recruitment events (MR-LF or MR-SO), (d) traffic jams, defined as instances where ants could not move at least one body length (1 cm) in 4 seconds, with at least one tandem pair or brood transporter is also stuck behind them. However, as per previous literature, they are capable of moving at the speed of ∼4cm/s and can easily pass through the focal area (15 cm) in 4 seconds.

### Agent-Based Simulation

The agent-based simulation (ABM) provided a virtual environment to explore the dynamics of colony transport by tandem running in *D. indicum* that cannot be examined experimentally. Technical details of ABM regarding an initial setup of system, update protocol and evolution of leader and follower during relocation is provided in the supplementary information (at point 3). A comprehensive description of the simulation protocol is presented in Fig. 1B. In the numerical simulation of the multi-lane condition, we modelled the space over which the ants move as a 2D array of cells. Along the *x*-axis (horizontal) are 152 cells, with the 0th and 151st cells representing the old and new nests, respectively. Along the *y*-axis (vertical), there are 5 cells at each *x*-position. Given that an adult *D. indicum* is typically 1 cm long and 0.5 cm wide, it occupies 2 cells along the *x*-axis and 1 cell along the *y*-axis. Therefore, we kept each 2D cell’s dimensions of 0.5 cm x 0.5 cm, and the path between the two nests is 75 cm long and 2.5 cm wide. In the numerical simulation of the one-lane condition (bottleneck), we constricted the 2D array in the middle such that from the 60th to the 89th cell along the *x*-axis, there is only one cell along the *y*-axis. This creates a constricted path that is 15 cm long, simulating the bottleneck in the experimental setup.

Our code updates the state of the system mimicking the experimental setup at hand after every one second. We mathematize the system so that various parameters, variables, and measurable outcomes can be quantified. For tagging ants’ states, we denote the state (at time *t* ∈ {0, 1, 2,*…* } second) of an ant, which is either a tandem leader or a brood carrier or a returning leader, with *L*_*i*_ where *i* can run from 0 to *n*_*s*_; and denote the state (at time *t* ∈ {0, 1, 2,*…*} second) of an ant, acting as a follower, by 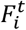 where *i* can run from *n*_*p*_ + 1 to *N* − 1. These states explicitly depend on certain variables and parameters as mathematically represented below:

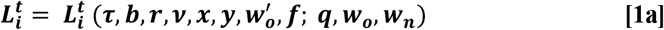

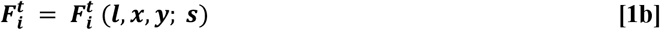

The symbols have following meanings:

***(τ)*:** It can take binary value, 1 or 0, depending on whether the leader is tandem leader or not.

***(b*):** It can take binary value, 1 or 0, depending on whether the leader is brood carrier or not.

**(*r*):** It can take binary value, 1 or 0, depending on whether the leader is returning leader or not.

***(ν*):** It denotes the number of completed tandem runs; hence, it is necessarily a non-negative integer.

**(*x*):** The cell-coordinate in *x*-direction—*x* ∈ {0, 1,*…*, 151}. Since an ant has been modeled to occupy 2 cells along *x*-axis, we adopt the convention that *x* specifies the front cell out of the two.

***(y*):** The cell-coordinate in *y*-direction—*y* ∈ {0, 1,*…*, 4} in the non-constricted part of the track and *y* ∈ {2} in the constricted part of the track.

(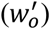): The time (in seconds) required by a returning leader to initiate a tandem run having reached the old nest. It is a random number chosen from the uniformly discretized interval [9, 26].

**(*f*):** It is the integer tag of the follower following the *i*^th^ tandem leader. It can take values from *n*_*p*_ + 1 to *N* − 1 when *τ* = 1. For the reason of book-keeping, when *b* = 1 or *r* = 1, *f* is set as −1.

**(*q*):** The probability that a leader ant gives up on becoming returning leader (i.e., does not come out of the new nest) after completing one tandem run.

(*w*_*o*_): This is the waiting time (in seconds) that a primary leader (ant with tag 1 to *n*_*p*_ ) takes to initiate her first tandem run from the old nest. The 0th ant is assigned *w*_*o*_ = 0; for others, it is a random number chosen from the uniformly discretized interval [0, 900].

(*w*_*n*_): This is the waiting time (in seconds) for a secondary leader (after reaching the new nest as a follower) to emerge out of the new nest as a returning leader. It is a random number chosen from the uniformly discretized interval [0, 900].

**(*l*):** It can take binary value, 1 or 0, depending on whether the follower is a lost one or not.

**(*s*):** It can take binary value, 1 or 0, depending on whether the follower is a secondary leader or not.

At the beginning of the simulation, *N* ants—ranging from 54 to 160, reflecting the range of colony sizes in our experiments—occupy the old nest. We categorized these ants as primary leaders (10%) and followers (90%). Secondary leaders constitute around 10% of the total ant population among the followers. Any ant acting as a leader (primary or secondary) also acts as a brood carrier with a probability of 0.1; otherwise, it tandem runs a follower. We consider the relocation complete when all ants from the old nest have reached the new nest.

Regarding ant movement, each ant moves with a constant speed—the exact value depends on the ant’s type (tandem leader, follower, brood transporter or returning leader)—along the *x*-axis. Along the *y*-axis, the ant performs a lazy random walk. We update all ant positions randomly and asynchronously to transition the entire system to its next state. No two ants occupy the same position; we model the effects of interactions between ants attempting to occupy the same cell based on experimental observations. The cells corresponding to the old and new nests are exceptions, as there is no limit on the number of ants occupying them. These interactions include creating lost ants and their subsequent recruitment, the switching over of followers, the turning back of returning leaders, and the emergence and resolution of jams.

We utilized experimental evidence to calibrate various simulation parameters associated with ant behaviour. To establish causal relationships between the observed phenomenon of traffic jams and their resolution, we conducted a comprehensive examination of the impact of each individual behaviour on the system dynamics. However, isolating and manipulating a single behaviour in the experimental framework is not feasible, as controlling the actions of individual ants in such a precise manner is impractical. The ABM overcomes this limitation by providing a platform for systematically varying the parameters that govern specific ant behaviour, allowing us to assess their isolated impact on jam resolution. For instance, to investigate the role of the switching-over behaviour, we can manipulate the corresponding parameter. By completely switching off this parameter, we can simulate a scenario where no switching-over occurs, thus predicting jam resolution times in its absence.

Furthermore, the model allows for systematically exploring other parameters and probabilities associated with underlying processes, providing a comprehensive understanding of the interplay between individual behaviour and collective outcomes. We iterated all simulations 25 times for each colony (*n* = 12) and maintained colony sizes at the mean size of the experimental ant colonies (87 ants).

### Statistical Analysis

We analyzed experimental data using Wilcoxon paired-sample tests as same colonies were relocated in control and manipulation experiments. Whereas, the ABM simulation data were analyzed by using Mann-Whitney *U* tests as at each simulation different agent function as the ant in a colony. Spearman’s correlation was used to measure any correlation between colony size and jam frequency. The Jam creator’s active contribution during experimental observation was analyzed between observed vs. expected using Chi-Square test. We set the statistical significance level at *P* < 0.05.

## Acknowledgements

We thank Basudev Ghosh for assistance with ant colonies. We acknowledge Debashish Chowdhury for discussions on stochastic models and Raghvendra Gadagkar for valuable comments on the manuscript. Rajbir Kaur and Udipta Chakraborty generously provided insights to improve readability of the final version of this article.

## Funding

Department of Science and Technology, India-INSPIRE fellowship 17IF0031 (MP).

Ministry of Education, India-Prime Minister’s Research Fellowship (JBD).

Science and Engineering Research Board, India-MTR/2021/000119 (SC)

Science and Engineering Research Board, India-EMR/2017/001457 (AS).

## Author contributions

Conceptualization: MP, SA

Methodology: MP, JBD, SC, SA

Data collection & analysis—experiment: MP

Agent based simulation & analysis: JBD, SC

Writing—original draft: MP, JBD

Writing—review & editing: SA, SC

Supervision: SA, SC

## Data and materials availability

All data are available in the main text or the supplementary materials. The model simulation code is available on GitHub [[link](https://github.com/joydasbairagya/Traffic-Jams-and-its-resolution-in-tandem-running-ants)].

## Supplementary video 1: (click here for video)

**Suppl video 1: Traffic Jams and its resolution in tandem-running *D. indicum* ants**. This video from our experiment showcases the dynamics of traffic flow in *D. indicum* ants during nest relocation along one and multi-lane path. It displays how jams are caused by leaders returning from the old nest, who interrupt the flow. After ants struggles in the jam for multiple seconds, mass orientation towards the new nest occurs with majority of ants aligning towards the new nest and clear the jam. Events of returning leaders actively recruiting nestmates on this congested path can also be seen. The traffic on the multilane path remains smooth even on high density.

