## Supplementary information 1 for "Ants get stuck in traffic jams and resolve them by making adaptive decisions"

##### This PDF file includes:

Supporting text
Figs. S1 to S11
Table S1
SI References

### Supporting Information Text

#### 1. Other potential jam mitigation strategies

Several potential strategies for mitigating traffic jams in *D. indicum* were explored, although their empirical support was limited. One hypothesis suggested that reducing the number of tandem leaders involved in relocation could decrease transport rates and, consequently, jam frequency. However, our observations revealed no significant difference in the involvement of tandem leaders between one-lane (20.54%) and multi-lane (18.07%) relocations, refuting this hypothesis. Another potential strategy involved the enforcement of traffic regulations by ants stationed at the entry and exit points of the constrained pathway. However, our observations did not support this hypothesis either. No ants consistently occupied these positions, nor did they exhibit any regulatory behaviors such as interfering with traffic or displaying aggression. Lastly, we considered the possibility of synchronized, unidirectional traffic flow as a solution to path constraints. However, analysis of relocation frames revealed no evidence of such coordination. Instead, the proportion of ants moving onward and those returning remained consistently balanced throughout the observed period (mean proportion = 0.54). While these potential strategies were not observed in our study, they may be employed by *D. indicum* under different ecological contexts or by other ant species facing similar challenges. Future research should explore the broader repertoire of traffic management strategies employed by ants in diverse environments and social contexts.

#### 2. Other phases of relocation dynamics

There are two phases of nest relocation that are not included in our agent-based modeling (ABM; detailed in the next section): discovery phase and latency phase. In the discovery phase, ants endeavour to discover a new nest from the moment the old nest is deemed uninhabitable. The time taken for the first discovery of the new nest is called the *discovery time*. During the latency phase, the ant colony initiates its first tandem run. The time between the discovery of new nest and the initiation of first tandem run, is known as the *latency time*. Statistically speaking (Wilcoxon paired-sample test, $P > 0.05$), neither the discovery time nor the latency time depends on whether the relocation path is one-lane or multi-lane. Specifically, we found that discovery time was $2.96 \pm 2.90$ minutes in one-lane vs. $4.68 \pm 3.13$ minutes in multi-lane, and the latency time was $6.59 \pm 6.04$ minutes in one-lane vs. $5.17 \pm 2.78$ minutes in multi-lane. As experimental observations indicate that these two phases are unaffected by path constriction, and furthermore, not obviously involved in jam phenomenon, we excluded them from our ABM.

#### 3. Technical details of ABM

**A. Initial setup of system.** In our numerical simulation of multi-lane experiments, the space over which ants move is modeled as a 2D array (FIG. S1 $(a)$): Along $x$-axis (horizontal, say), there are 152 cells—the 0th and the 151th cells represent the old and the new nests, respectively—at each $y$-position. Along vertical $y$-axis (perpendicular to the $x$-axis), there are 5 cells at each $x$-position. One adult ant of the species *D. indicum* is typically 1 cm long and 0.5 cm wide and is modeled to occupy 2 cells along $x$-axis and 1 cell along $y$-axis. This means each 2D cell has dimension 0.5 cm $\times$ 0.5 cm. While there is no limit on how many ants can occupy the cells corresponding to the old and the new nests, two ants are not allowed to simultaneously exist on any other cell. In the numerical simulation of one-lane setup (see FIG. S1 $(b)$), the 2D array is constricted in the middle so that from 60th cell to 89th cell along $x$-axis, there is only one cell along the $y$-axis.

To begin with, $N$ (ranging from 54 to 160) number of ants occupy the old nest. These ants are categorize in the following way: primary leaders (10%) and followers (90%). Among the followers, there are secondary leaders who constitute 10% of the total ant population. Moreover, any leader (primary and secondary), going from old to new nest, may act as brood carrier with probability, $p_{\rm bc} = 0.1$; otherwise, she guides a follower.We have assumed there are a lot of broods and transport of the colony is deemed completed when all the ants from the old nest reach the new nest, irrespective of the number of broods remaining in the old nest.

It is convenient to tag each ant in the old nest using a unique integer between 0 to $N-1$: From 0 to $n_p$ (the integer nearest to one less than 10% of $N$) are reserved for primary leaders, next 10% integers—$n_p+1$ to $n_s$—are reserved for secondary leaders (a type of followers), and the rest 80% integers (i.e., $n_s+1$ to $N-1$) are assigned to the followers.

**B. Update protocol.** Our code updates the state of the system mimicking the experimental setup at hand after every one second. Let us mathematize the system so that various parameters, variables, and measurable outcomes can be quantified.

***B.1. Tagging ants' states.*** To this end, we denote the state (at time $t \in \{0,1,2,\cdots\}$ second) of an ant, which is either a tandem leader or a brood carrier or a returning leader, with $L_i^t$ where $i$ can run from 0 to $n_s$; and denote the state (at time $t \in \{0,1,2,\cdots\}$ second) of an ant, acting as a follower, by $F_i^t$ where $i$ can run from $n_p+1$ to $N-1$. These states explicitly depend on certain variables and parameters as mathematically represented below:

$$L_i^t = L_i^t(\tau, b, r, \nu, x, y, w_o', f; q, w_o, w_n), \qquad [1a]$$

$$F_i^t = F_i^t(l, x, y; s). \qquad [1b]$$

Here, the symbols have following meanings:

1. $\tau$: It can take binary value, 1 or 0, depending on whether the leader is tandem leader or not.

2. $b$: It can take binary value, 1 or 0, depending on whether the leader is brood carrier or not.

3. $r$: It can take binary value, 1 or 0, depending on whether the leader is returning leader or not.

4. $\nu$: It denotes the number of completed tandem runs; hence, it is necessarily a non-negative integer.

5. $x$: The cell-coordinate in $x$-direction—$x \in \{0, 1, \cdots, 151\}$. Since an ant has been modeled to occupy 2 cells along $x$-axis, we adopt the convention that $x$ specifies the front cell out of the two.

6. $y$: The cell-coordinate in $y$-direction—$y \in \{0, 1, \cdots, 4\}$ in the non-constricted part of the track and $y \in \{2\}$ in the constricted part of the track.

7. $w'_o$: The time (in seconds) required by a returning leader to initiate a tandem run having reached the old nest. It is a random number chosen from the uniformly discretized interval $[9, 26]$.

8. $f$: It is the integer tag of the follower following the $i$th tandem leader. It can take values from $n_p + 1$ to $N - 1$ when $\tau = 1$. For the reason of bookkeeping, when $b = 1$ or $r = 1$, $f$ is set as $-1$.

9. $q$: The probability that a leader ant gives up on becoming returning leader (i.e., does not come out of the new nest) after completing one tandem run. However, at any given instant before the transport is complete, all leaders are not allowed to be quitters together.

10. $w_o$: This is the waiting time (in seconds) that a primary leader (ant with tag 1 to $n_p$) takes to initiate her first tandem run from the old nest. The 0th ant is assigned $w_0 = 0$; for others, it is a random number chosen from the uniformly discretized interval $[0, 900]$.

11. $w_n$: This is the waiting time (in seconds) for a secondary leader (after reaching the new nest as a follower) to emerge out of the new nest as a returning leader. It is a random number chosen from the uniformly discretized interval $[0, 900]$.

12. $l$: It can take binary value, 1 or 0, depending on whether the follower is a lost one or not.

13. $s$: It can take binary value, 1 or 0, depending on whether the follower is a secondary leader or not.

***B.2. Evolution of $L_i^t$ and $F_i^t$.*** Our numerical code is all about how $L_i^t$ and $F_i^t$ evolve as time, $t$, flows.

**Specifying $L_i^0$ and $F_i^0$**: At time $t = 0$, we set the initial condition as

$$L_0^0(\tau = 1, b = 0, r = 0, \nu = 0, x = 0, y = y_0, w'_o = 0, f = \eta_0; q = 0.001, w_o = 0, w_n = 0); \quad [2a]$$

$$L_i^0(\tau = H(p - 0.1), b = 1 - H(p - 0.1), r = 0, \nu = 0, x = 0, y = y_0, w'_o = 0, f = \eta_i \tau - b; q = 0.001, w_o = w_{oi}, w_n = 0), \quad \forall i \in \{1, \cdots, n_p\}; \quad [2b]$$

$$F_i^0(l = 0, x = 0, y = y_0; s = 1 - H(i - n_s - 1)), \forall i \in \{n_p + 1, \cdots, N - 1\}. \quad [2c]$$

Here, $y_0$ is a number randomly chosen from set $\{0, 1, 2, 3, 4\}$ and $\eta_i \in \{n_p + 1, \cdots, N - 1\}$ (for each $i \in \{0, \cdots, n_p\}$) is unbiasedly chosen random integer. $H$ denotes the Heaviside step function where $p$ is a random number picked unbiasdly from the interval $[0, 1]$. This incorporates the fact that, except during the first tandem run, any leader in the old nest becomes brood transporter/carrier with a probability $p_{\text{bc}} = 0.1$; otherwise she carries a follower. Also, $w_{oi}$ (for each $i \in \{1, \cdots, n_p\}$) is a realization of the random number $w_o$ deciding when the $i$th leader will come out of the old nest.

When an ant is outside the nests, only four distinct basic movements are allowed in the absence of any other ant on the track:

1. Tandem leader: A leader carrying a follower moves towards right (new nest). She moves with a speed $v_T$ along $x$-axis. It means in our simulations, she moves 8 cells/second along $x$-direction. She can simultaneously move along $y$-direction, i.e., we allow her to stray vertically while moving horizontally. The probability of moving up at each $x$-position is $\alpha$; the probability of moving down is same. Obviously at the lowermost and uppermost cells, the ant is constrained to stray with probability $\alpha$ away from the boundary along $y$-direction.

2. Brood transporter: A leader carrying brood moves exactly as the tandem pair except that her speed along $x$-axis is $v_B$.

3. Returning leader: A returning leader—a leader moving towards old nest (left) to seek a new follower—also moves exactly as the tandem pair except that her speed along $x$-axis is $v_R$.

4. Lost follower: A follower who loses a leader is modeled to perform an unbiased random walk along $x$-axis. In the $y$-direction, she moves like the other aforementioned ants.

Now let us see how ants' movements update the state of system when there is no interaction among them. Of course, when more than one ants are on the track, they may interact leading to some modified movements which will be discussed later.

**Going from $t$ to $t+1$**: Naturally, we may denote the state of the entire system of ants collectively as $(\{L_i^t\}, \{F_i^t\})$. Given $(\{L_i^t\}, \{F_i^t\})$, the code iteratively calculates $(\{L_i^{t+1}\}, \{F_i^{t+1}\})$ till all the ants are transferred from the old nest to the new nest. To this end, the tags, 0 to $n_p$, of the leaders are sorted randomly as an array of numbers. Subsequently, the numbers are chosen sequentially and the corresponding leaders' states are updated asynchronously: After one ant's coordinates are updated by one cell, the next ant is chosen for update; and this process continues till all the ants' coordinates are updated by at most one cell. Then, the tags, 0 to $n_p$, of the leaders are re-sorted randomly, and the above process repeats till one second is over. As far as the followers' states are concerned, only those are updated that are associated with the followers picked by the leaders.

Let us understand the exact updation process of a single ant's coordinates. For illustration, consider a leader during the time duration when $t$ changes from 0 to 1. During update of $i$th leader's state (assuming $t = 0 \geq w_{oi}$; obviously satisfied by 0th leader at least), the $x$-coordinate and the $y$-coordinate of the leader are evolved as follows: $x$-coordinate explicitly increases cell-by-cell for $2v_T$ cells (if $\tau = 1$) or $2v_B$ cells (if $b = 1$), whereas $y$-coordinate at each $x$-coordinate increases or decreases by one cell with probability $\alpha$. (Of course, at the boundaries, the $y$-coordinate is updated such that with probability $\alpha$, the ant moves away from the boundary.) While at a mean-field level it doesn't matter whether $x$-coordinate or $y$-coordinate is updated first; for concreteness, we have always updated $y$-coordinate first. If the $i$th leader, chosen for update, is such that $t = 0 < w_{oi}$, then her state remains unchanged after one second. During the update of $L_i$ (with $t = 0 \geq w_{oi}$ and $\tau = 1$), the state of follower with tag $\eta_i$ is updated simultaneously in such a manner that $x$-coordinate of the follower is two less than that of her leader. The $y$-coordinate of the follower is same as that of her leader.

For further clarity of the updation process, consider a hypothetical case of a colony with only two leaders—one of them is tandem leader and the other is brood transporter at time $t$—such that $w_o = 0$. We assume, for simplicity, that during the process, one ant doesn't obstruct the path of the other ant at any step of the process. At first, we chose one of them randomly and update its $y$-coordinate by one cell and then update its $x$-coordinate by one cell. Thereafter, we similarly update the second ant's $y$- and $x$-coordinates by one cell each. We repeat the process until leaders complete their respective movements in one second—the tandem leader moves $2v_T$ cells in $x$-direction, whereas the brood transporter moves $2v_B$ steps. Thus, finally, we deem the state of this colony updated from its configuration at $t$ to that at $t+1$.

Note that outside the nests, a tandem pair occupies four adjacent cells on the track along $x$-axis. Since the updates (over each second) are asynchronous, when another tandem pair comes out of the old nest during its turn of getting updated, it may find the earlier updated tandem pair on its way (assuming that the $y$-coordinate is same as that of the former tandem pair). Since two ants cannot occupy same cell, the latter tandem pair halts directly behind the follower of the tandem pair in front; however, if it strays along $y$-axis (allowed with probability $2\alpha$), it may further its location along $x$-axis. Since, it can equivalently stray along $y$-direction with probability $2\alpha$ at each remaining steps out of total $2v_T$ steps. In our code, we similarly model the interactions involving brood-carriers keeping in mind that in such cases, $v_B$ (and not $v_T$) decides total number of cells.

Finally, as per the update rules delineated in the two immediately preceding paragraphs, the code generates the system's state $(\{L_i^1\}, \{F_i^1\})$ from its initial state $(\{L_i^0\}, \{F_i^0\})$. This process continues in the same manner with no new phenomenon till the new nest is reached by the ants: In our numerical simulations, a leader of the tandem pair is said to have completed the transport of her follower in the new nest when the entire body of the leader enters the new nest; and we assign $x$-coordinate as 151 to both the leader and her follower.

**Acts inside new nest**: Suppose that at time $t$, $i$th leader enters the new nest. If $\tau = 1$ for the leader, then $\nu$ increases by one unit since one successful tandem run has been accomplished; furthermore, her $f$ variable's value changes from $\eta_i$ to $-1$. The leader can now decide whether to come out of the nest to initiate her next tandem run: She may decide not to come out with probability $q$; in this case, $L_i$ is not updated anymore. In the case (occuring with probability $1-q$) $L_i$ is to be updated, the $r$ variable corresponding to the leader takes value 1; variables $\tau$ (if the leader was tandem leader) or $b$ (if the leader was brood carrier) are reset to 0. During the sequential updating, when this leader's turn comes, her $x$-coordinate explicitly decreases step-by-step for $2v_R$ steps, whereas $y$-coordinate at each $x$-coordinate increases (decreases) by one unit with probability $\alpha$ (with usual care taken at the boundaries); thus, $L_i^t$ updates to $L_i^{t+1}$ with $r = 1$. As returning leaders start their marches on the track, there is the possibility of their (returning leader interacts with another returning leader) interaction during the sequential updating as is the case with tandem (or brood-carrier) leaders detailed earlier. The update rule in such interactions is same as that described above for the tandem leaders.

We also have to keep in mind that if the leader is a tandem leader, then she guides a follower, tagged $\eta_i$, with it. If the corresponding $s = 0$, then $F_{\eta_i}^t$ is not updated anymore. If, however, $s = 1$, $F_{\eta_i}^t$ is renamed as $L_{\eta_i}^t$ (with $\tau = b = r = 0$), i.e, the follower has become a leader at rest in the new nest; parameter $w_n$ for this leader (with $x = 151$) changes from 0 to $w_{n\eta_i}$—a number randomly chosen from interval [0,15]. If $w_{n\eta_i} \leq 1$, then $L_{\eta_i}^t$ updates to $L_{\eta_i}^{t+1}$ with $r = 1$, $\tau = b = 0$, $\nu = 0$, $f = -1$, $w_o = 0$ and $w_n = w_{n\eta_i}$. Its subsequent evolution is similar to that of the $i$th leader detailed above. If $w_{n\eta_i} > 1$, then the variables and parameters inside $L_{\eta_i}^t$ remain unchanged at $t+1$ and the follower-turned leader remains at rest in the new nest until in the future series of sequential updates at a time $t+m$ with the least $m$ (a postive integer) comes such that $w_{n\eta_i} \leq m$. Then the leader at rest will come out of the nest with $L_{\eta_i}^t$ updated to $L_{\eta_i}^{t+m}$ with $r = 1$, $\tau = b = 0$, $\nu = 0$, $f = -1$, $w_o = 0$ and $w_n = w_{n\eta_i}$.

**Acts inside old nest**: The returning leader tries to get back to the old nest to fetch more followers and broods. When the entire body of the returning leader—let's call it $j$th leader for convenience—enters the old nest at $t = t'$ (say), the variables and parameters corresponding to her state $L_j^{t'}$ are set as: $r = 0$, $\tau = H(p - 0.1)$, $b = 1 - H(p - 0.1)$, $\nu$ =some number depending on the history, $x = 0$, $y = 2$, $w'_o = w'_{oj}$, $f = \eta_j \tau - b$, $q = 0.001$, $w_o = 0$, and $w_n = 0$. Here, $w'_{oj}$ is a random number chosen from uniformly discretized interval $[9, 26]$. The $j$th leader remains at rest in the old nest until in the future series of sequential updates, a time $t + m$ with the least $m$ (a postive integer) comes such that $w'_{oj} \leq m$. Then the leader will come out of the nest.

**Interactions between ants**:

More interestingly, when more than one ant are on the track traveling in opposite directions, movement of the ants should be constrained further. While traveling in any allowed direction if an ant finds another ant occupying a cell on her path, then it is not allowed to simultaneously occupy that cell or hop over the occupied cell. However, she can perform other allowed movements with specified rates. As observed in the experiments, interactions between ants lead to some important phenomena that we incorporate in our numerical simulations:

1. Constrained vertical movement during head-on encounter: When a tandem pair or a brood transporter meets a returning leader head-on, we say that an *encounter* has occurred. The tandem pair or the brood transporter receives a call of the returning leader and its straying along $y$-direction: Effectively, it means that $\alpha$ is reduced by a factor $\beta$ till the encounter is sustained in an immobile state.

2. Turning of returning leader in an encounter: During an encounter, the returning leader waits for $T_r$ seconds starting from the instant of encounter; and then, with a probability rate $\rho$ per step she may turn back towards the new nest. Note that an encounter need not happen at integral times: They mostly occur during the updates and recorded after every one second. In this context, we point out that if during an encounter, the tandem pair (or brood transporter) and the returning leader remain immobile for two seconds or more, we say that an interruption has occurred. Therefore, if $T_r \geq 2$ seconds, then one may say that the turning of returning leaders occurs only during interruptions. Jumping ahead of ourselves (see point 5 below), during jams, $T_r$ and $\rho$—now denoted by $T_r^J$ and $\rho^J$, respectively—may be chosen to be different. A special situation should be mentioned here: A returning leader with location $(x, y)$=$(91, y \in \{0, 1, 3, 4\})$, although technically is not in an encounter situation, is also allowed to turn back as if it has faced an encounter.

3. Follower leaving leader in an encounter: After waiting for $T_l$ time in an encounter between a tandem pair and a returning leader, the follower in the tandem pair may decide to leave her leader. Its future now follows either of the two following events that are taken to occur, respectively, with probabilities $p_{\text{so}}$ and $1 - p_{\text{so}}$:

   a. Switching-over: The follower in the tandem pair leaves her leader to accept the invitation of the returning leader with a view to forming a new tandem pair that heads towards the new nest. (The leader of old tandem pair heads back towards the old nest.) This phenomenon is termed *switching over*. Let us say that the initial tandem pair had $i$th leader and $\eta_i$th follower, and the returning leader was the $j$th ant. At switching over, $i$th leader's state is reset such that $\tau = 0$ but $r = 1$ and $f = -1$, whereas $j$th leader's state is reset such that $r = 0$ but $\tau = 1$ and $f = \eta_i$.

   b. Creation of lost ants: The follower looses the contact with her leader and becomes a leaderless follower—appositely termed a *lost ant*—and wanders around performing a 2D random walk. (The leader of the tandem pair keeps going towards the new nest.) Let us say that the initial tandem pair had $i$th leader and $\eta_i$th follower, and the returning leader was the $j$th ant. At the event of losing the leader, $i$th leader's state is reset such that $\tau = 0$ and $f = -1$, whereas $\eta_i$th follower's state is reset such that $l = 1$.

   Sometimes a tandem pair is unable to move in the forward direction and waits behind of an ant, i.e., anterior of the tandem pair's leader touches posterior of another ant (e.g., follower in a tandem pair, lost ant, brood transporter, returning leader who is moving towards the new nest). By definition, this situation is not counted as an encounter since their interactions are not head-on. While in such interactions, switching over is not possible as no returning leader is trying to recruit the follower of the tandem pair, there is possibility of creation of lost ant: If the tandem pair waits more than $T_l$ time, the follower may leave her leader (say, $i$th leader) with probability $\lambda$ per step and becomes a lost ant. In this event, $i$th leader's state is reset such that $\tau = 0$ and $f = -1$, whereas $\eta_i$th follower's state is reset such that $l = 1$. Additionally, we include the possibility of creation of lost ant when no obstacle is present in front of the tandem pair: The follower may become lost with probability $\lambda$ per step after successfully follow a leader more than $T'_l$ time.

4. Mid-way tandem recruitment: There are two ways of recruitment in the midway—switching over and recruitment of lost ants. The former one is as discussed above and the latter one is due to the interaction between a follower-less leader and a lost ant. (An event, where a leader finds back her own lost ant, is not considered as a midway-recuirtment event.) If a follower-less leader's anterior touches a lost ant (recall that every ant spans two adjacent horizontal cells), i.e., along $x$-axis they do not have any vacant cell in between, then they form a tandem pair with probability $p_{\text{lfr}}$ per step and both head towards the new nest. Let us say that the $j$th leader meets $i$th lost follower, then $j$th leader's state is reset such that $\tau = 1$ but $r = 0$ and $f = i$, whereas $i$th follower's state is reset such that $l = 0$.

5. Jam: *Jam* is the phenomenon that is central to our investigation and in our simulations it emerges automatically. In our simulations, for concreteness and simplicity, a jam is said to have occurred, if during an encounter, another tandem pair (or brood transporter) settles directly behind the tandem pair (or the brood transporter) with no empty cell in between, and they all remain immobile for at least four seconds. It should be kept in mind that in real experiments, complete

immobilisation of an ant does not happen: An ant is defined to be immobile if she cannot move a distance equivalent to her one body length. In passing, we remark that results reported here remain statistically the same if we weaken the definition of interruption and jam. In the modified definition, a head-on encounter is regarded as an interruption if two leaders (not necessarily involving a follower)—one heading towards the new nest and the other towards the old nest—remain immobile for at least 2 seconds. Similarly, a jam could be said to have occurred when the interruption is prolonged, and another leader (not necessarily guiding a follower) heading towards the new nest has to wait still for at least 4s behind the interrupting pair.

Naturally, every jam is an interruption but not vice versa. One of our main quantity of concern is 'jam resolution time' that is the time taken after the aforementioned four seconds to resolved the jam.

We take this opportunity to point out which kinds of jams our simulations exactly deal with. We know that while carrying a pupa with their mandibles, brood transporters face hindrance to their movements, especially on the one-lane path. Despite creating 27% of all observed jams due to transportation inefficiencies in the one-lane path, brood transporters seemingly play a negligible role in jam resolution. As our primary focus is on jam resolution, we have concentrated solely on jams caused by lost followers and returning leaders to effectively pinpoint the role of returning leaders in mitigating congestion.

6. Returning-returning leaders' interaction: When a returning leader turns during the jam resolution, she may find another adjacent returning leader (say, 2nd returning leader) head-on. For jam resolution, the 2nd returning leader must also turn back. The protocol adopted for the 2nd returning leader's turning is similar: The 2nd returning leader waits for at least $T_r$ seconds starting from the instant of touching the posterior of the 1st returning leader till the latter turns and faces it head on; subsequently, with a probability rate $\rho$ per step she may turn back towards the new nest. Of course, a series of similar interactions, like between the 2nd returning leader and a 3rd returning leader, is possible. In all such interactions in our simulations, the returning ants follow the same protocol for turning. We term this protocol *coordinated turning*.

| **Parameters** | **Meaning of the parameters** | **Values used in simulations** |
|---|---|---|
| $N$ | Size of ant colony | $54-160$ |
| $\frac{n_p+1}{N}\times 100$ | Percentage of primary leaders | 10% |
| $\frac{n_s-n_p}{N}\times 100$ | Percentage of secondary leaders | 10% |
| $p_{bc}$ | Probability of becoming a brood transporter | 0.1 |
| $\alpha$ | Probability of moving up or down along $y$-direction | 0.025 |
| $\beta$ | Factor of reduction in $\alpha$ during encounter | 0.3 |
| $v_T$ | Velocity along $x$-direction of tandem pair | 4 cm/second |
| $v_B$ | Velocity of a brood carrier | 4.5 cm/second |
| $v_R$ | Velocity of a returning leader | 6.5 cm/second |
| $T_l$ | Time taken for becoming lost ant for an immobile tandem pair | 4 seconds (non-jam case) or 9 seconds (jam case) |
| $T_l'$ | Time taken for becoming lost ant for a mobile tandem pair | 4 seconds |
| $p_{so}$ | Probability of switching over in an encounter | 0.3 |
| $\lambda$ | Probability of becoming lost | 0.001/step |
| $T_r$ | Time taken for turning towards new nest in non-jam situation | 4 seconds (non-jam case) or 9 seconds (jam case) |
| $\rho$ | Probability of turning back in an interaction | 0.02/step (non-jam case) or 0.007/step (jam case) |
| $p_{lfr}$ | Probaility of making a tandem pair with a lost ant | 1 |
| $q$ | Probability of quit the role of a leader | 0.001/step |
| $w_o'$ | Time required to initiate a tandem run by a returning leader | [9,26]second uniformly picked |
| $w_o$ | Time required to initiate first tandem run | [0,15]min uniformly picked |
| $w_n$ | Time required for secondary leader to become returning leader | [0,15]min uniformly picked |

**Table S1. List of some parameters used in the simulations.**

**C. Choice of parameter values.** We note that there are many parameters (see Table S1) involved in the simulations: While some are fixed using the experimental results, some parameters are free. Let us now list here the quantities that we have used in the numerical simulations so as to benchmark our code against experiments, to understand the experimental results, and to make predictions:

1. $N$, $n_p$ and $n_s$: Their values used in the simulations correspond to what have been observed and documented in the experiments under consideration (1).

2. $v_T$, $v_B$ and $v_R$: These values have been taken from observations documented in literature (2).

3. $w_o'$: It has been obtained from direct observations (2).

4. $\alpha$: From observations (2, 3), we know the *path efficiency*—the average ratio of displacement to distance of an ant traveling from old nest to new nest—is around 80%. In simulation, we numerically calculate the path efficiency for various values of $\alpha$ in the 5-lane setup. Arguably, the path efficiency in the simulations should be even more as the breadth to length ratio is 1 : 30 which is much less than 2 : 3 used in the aforementioned experiments. Naturally, the path efficiency should depend on $\alpha$; consequently, by trail-and-error, we choose $\alpha = 0.05$ for which the path efficiency is 90%. While there is a bit of arbitrariness involved in this choice, the choice is further validated by the fact that distribution of total numbers of interruptions seen in direct observations and numerical simulations are statistically identical—à la the Mann–Whitney U test—in both one-lane and multi-lane cases [FIG. S2. $(a)$ and S3. $(a)(P \geq 0.05)$]. This comparison makes sense because the movement along $y$-direction (characterized by $\alpha$) is naturally correlated with the number of interruptions: the more frequent movements along $y$-axis are, the less frequent interruptions are.

5. $\beta$: It is a free parameter (factor) by which $\alpha$ is reduced during the duration of an encounter when the constituent ants are in an immobile state.

6. $T_l$: $T_l$ is a free parameter; however, observations suggest that it is of the order of 4 seconds, and that is what we have used in our simulations.

7. $p_{so}$ and $\lambda$: These are free parameters in our simulations. We have performed a trial-and-error fixing of the values of $p_{so}$ and $\lambda$ by making sure that the fraction of total tandem runs exhibiting switching over and the fraction of total tandem runs involving creation of lost ants match with the respective fractions obtained in the experiments. See Figs. S2 $(d)$ and $(e)$, and Figs. S3. $(d)$ and $(e)$ which validate that $p_{\text{so}} = 0.3$ and $\lambda = 0.001$ are reasonable choices.

8. $T_r$ and $T_r^J$: $T_r$ is also a free parameter. In the encounters that are not jams, observations suggest that it is of the order of 4 seconds, and that is what we have used in our simulations. In the case of jam, the returning leader takes more time to turn towards new nest. This is because, in a jam, more than one tandem pairs wait allowing the returning leader to invite more than one follower to leave their respective leaders. We have done some trail-and-error to fix $T_r^J = 9$ seconds such that it is not only more than 4 seconds, but also results in a distribution of jam resolution times from numerical simulations such that the distribution is statistically same as the one found from the experiments (see FIG. S4).

9. $\rho$ and $\rho^J$: These free parameters have been fixed to $\rho = 0.02$ and $\rho^J = 0.007$ so as to generate a distribution of transport times that mimicks the distribution one gets from the experimental data (see FIG. S2 $(c)$ and FIG. S3 $(c)$).

10. $p_{lfr}$: While this is a free parameter, it makes sense in real scenario to keep its value as one because, almost with full certainity, any lost ant found by a returning leader is carried to the new nest.

11. $q$, $w_o$ and $w_n$: These three important free parameters predominantly decide the percentage of recruited leaders at any point of time. Hence, these have been fixed via trial-and-error in such a way that plot of the percentage of recruited leaders vs. the percentage of transport time elapsed matches qualitatively with that obtained from the direct experimental observations (1) (see FIG. S5).

**D. Results.** Having fixed the parameters in line with the experimental results in hand, we first simulate the experiments performed on 12 colonies of sizes 54, 56, 57, 68, 73, 76, 79, 90, 92, 110, 130 and 160, by taking 25 realizations of each colony. We specifically look for the statistical equivalence between the simulated and the experimental results on interruption, jam, transport time, switching over phenomenon, and lost ant recruitment in both the multi-lane and the one-lane setup: As seen in Figs. S2 and S3, our numerical code mimics the behaviour of the ants very closely. The justified confidence in our code encourages us to investigate how important the mid-way recruitment is for resolution of the jam events. For this purpose, we must work with a fixed-sized colony; hence, we take the size to be 87—average of the aforementioned 12 sizes—for our investigation as reported below. Moreover, to check for the robustness of our conclusions, we repeat the investigations for the smallest-sized (54) and the largest-sized (160) colonies as well.

We check the importance of the midway recruitment in two ways—midway recruitment related to lost ants and switching overs:

- Lost ant recruitment: We examine the importance of it by continuously decreasing the probability, $p_{lfr}$—something very hard, if not impossible, to control in experiments—of taking a lost ant by (mostly) a returning leader. It should be reiterated that we work under the assumption that $p_{\text{lfr}} = 1$ and $p_{\text{so}} = 0.3$ in the experimental (real-life) scenario. Now, as we decrease $p_{lfr}$ in simulations, the number of jams increases (see FIG. S6A(i)). However, we find the time distribution of jam resolutions corresponding to the experiment is qualitatively same as that found in simulations with either $p_{\text{lfr}} = 1$ or $p_{\text{lfr}} = 10^{-2}$ (see FIG. S6A(ii)). Furthermore, it may be pointed out that $p_{lfr}$ has direct effect on the transport time: As we decrease $p_{lfr}$ in simulations, the total transport time increases [see FIG. S6 A(iii)]. This is simply because the relocation of the entire colony is not readily possible if a lost ant may not be picked and taken by a leader ant to the new nest; after all, the only other way a sole lost ant can reach the new nest is by doing random walk which can take enormously long time, on average.

- Switching over: Similarly, in order to examine the role of switching over, we continuously decrease the probability of switching over, $p_{so}$, and keep $p_{\rm lfr} = 1$. Switching over plays a significant role in the creation of jams—a decrease in $p_{so}$ leads to a reduction in jams. As seen in Fig. S6B(i), a lower switching-over probability ($p_{so}$) results in fewer jams. A reason behind this may be as follows. Recall that, by definition, a jam occurs when one tandem pair waits (for four seconds) directly behind another one due to an encounter faced by the latter. However, when a lost ant is created (which happens with probability $1 - p_{so}$), the tandem leader becomes follower-less, and the lost follower remains just behind the leader (in simulations; in real experiments, they may orient themselves away from the leader)—This situation does not qualify as a jam. Thus, as $p_{so}$ decreases and the probability of lost ant creation increases, the number of true jams correspondingly decreases. Nevertheless, it seems the switching over phenomenon is helpful in resolving jam—increment in switching over probability effects resolution of jams [see FIG. S6 B (ii)]; consequently, the total transport time decreases as switching over increases [see FIG. S6 B (iii)].

It seems that the two midway recruitment processes affect the jams in two different ways: Recruitment of lost ants helps to avoid jams on narrow path and if a jam occurs, then switching over mechanism helps to resolve it quicker.

What, however, undoubtedly plays a crucial role in jams and its resolutions are $T_r$ and $\rho$. We may recall that during an encounter or a jam, the returning leader waits for $T_r$ (or $T_r^J$) seconds starting from the instant of encounter; and then, with a probability rate $\rho$ (or $\rho^J$) per step, it may turn back towards the new nest. To see their roles explicitly, first we continuously vary the value of $\rho$ corresponding to non-jam cases and keep the value of $\rho^J$ corresponding to jam cases fixed at 0.007. Now, as $\rho$ decreases, the propensity of an interruption becoming a jam increases [see FIG. S6 D (i)]. Although, it has no role in jam resolution since the probability of turning back during a jam ($\rho^J$) has been kept fixed [see FIG. S6 D (ii)]. As the decrement of $\rho$ delays the resolution of every encounter, the transport time should increase as seen in FIG. S6 D (iii).

Now, let us see the role of $\rho^J$. With a view to examining its role, we fix the value of $\rho$ to 0.02 while continuously changing the value of $\rho^J$. As expected, the larger is the value of $\rho^J$, the more is the chance of jam resolutions [see FIG. S7 B (ii)] and hence the less should be the transport time [see FIG. S7 B (iii)]. However, it should not have any effect on jam creation since, by construction, this probability is an effect of jam and not the cause [see FIG. S7 B (i)]. If we focus on the plot of percentage of jam resolutions versus the time taken for jam resolution in FIG. S7 B (ii), we note that the height of peak of the curve is dependent on the strength of $\rho^J$: As is intuitively obvious, the less the $\rho^J$, the lower should be the peak; this is exactly what we witness in simulations as exhibited in FIG. S7 B (ii).

Next let's see how do the parameters $T_r$ and $T_r^J$ regulate the time in which most of the jams resolve. If we increase the value of $T_r^J$ (corresponding to jam situation) by $\Delta T_r^J$ (while keeping the value of $T_r$ in the non-jam cases unchanged), then resolution time of a single jam should also increase. As expected, in our simulations, when we choose $\Delta T_r^J$ as 24 seconds, i.e., $T_r^J$ as 33 seconds, the maximum number of jams resolve around 33 seconds [see FIG. S8 B (ii)]. Clearly, the position of the peak, in the plot of percentage of jams versus the time required to resolve jams, predominantly depends on the value of minimum waiting time before turning back ($T_r^J$) after being stuck in a jam. Moreover, since resolution time of a single jam increases by same amount by which we increase $T_r^J$, in simulations, we do observe that the transport time increases [see FIG. S8 B (ii)] slightly. As may be guessed, by definition, changing $T_r^J$ should not has any effect on creation of jams; hence, the number of jams does not depend on the value of $T_r^J$ (see FIG. S8 B (i) for validation). However if we increase both $T_r$ and $T_r^J$ by $\Delta T_r$, as expected, the number of jams increases (see FIG. S6 C (i)). However, the distribution of jam resolving times depend only on $T_r^J$. Hence, for $\Delta T_r = 24$ (say), the distribution remains unchanged (compare FIG. S8 B (ii) with FIG. S6 C (ii). It should be noted that on increasing $T_r$ and $T_r^J$, every interruption that has been resolved by the turning back of returning leaders, should take more time to resolve. Ergo, the total transport time should increase more rapidly with increment $\Delta T_r$ (in both $T_r$ and $T_r^J$) than with increment $\Delta T_r^J$ (only in $T_r^J$) [compare FIG. S6 C (iii) and FIG. S8 B (iii).]

Lastly, we would like to know how much the 'coordinated turning' protocol is important to realize the aforementioned effects of $\rho$. To this end, we may adopt a different protocol, termed *uncoordinated turning*, that differs from the coordinated turning in the following way: Here, the 2nd returning leader waits for at least $T_r$ seconds starting from the instant of touching the posterior of the 1st returning leader; subsequently, with a probability rate $\rho$ per step she may turn back towards the new nest. Most importantly, in this protocol, the turning of the 2nd returning leader is not contingent on its becoming head-on with the 1st returning leader. Here, using both the protocols, we simulate the experiments performed on 12 colonies of sizes 54, 56, 57, 68, 73, 76, 79, 90, 92, 110, 130 and 160, by taking 25 realizations of each colony. The post-processed results are depicted in FIG. S9. It may be observed that the transport time, the number of interruptions, the number of jams and the jam resolution time are statistically same in both the protocols. Number of midway recruitments and number of switching overs are slightly more in the case of coordinated turning. This is simply because in coordinated turning protocol, in comparison with the uncoordinated one, there are chances of more encounters and hence, more chances of switching over phenomenon and creation of lost ants.

Finally, we mention that aforementioned conclusions about the role of $p_{lfr}$, $p_{so}$, $\rho$, $\rho^J$, and $T_r$ remain qualitatively unchanged even if larger or smaller than the average colony size (87) is considered in the simulations, see e.g., FIGs. S10, S11, S8, S7.

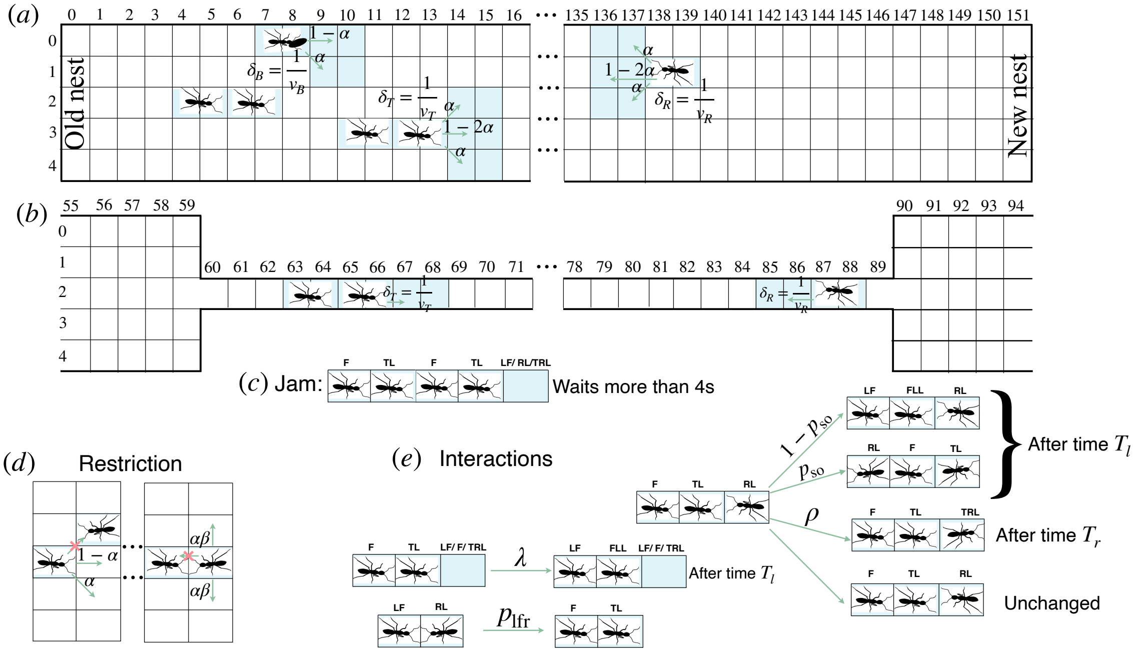


**Fig. S1.** Protocol used in the simulations: (a) and (b) depict the ants' movement during multi-lane and one-lane experiments. Green arrows show the possible y-directional movements with probability. Along the x-direction, a brood-transporter, a tandem pair and a returning leader move $\delta_B = 1/v_B$, $\delta_T = 1/v_T$ and $\delta_R = 1/v_R$ steps in one second, respectively. Subfigure (c) shows the definition of a jam, the letters F, TL, LF, RL and TRL represent follower, tandem leader, lost a follower, returning leader and turning back returning leader, respectively. Subfigure (d) is the pictorial representation of the fact that two individuals can not simultaneously occupy the same cell, and the tendency to remain longer in the same position if an ant finds another nest-mate is also introduced in the code by restricting the y-directional movement by a factor $\beta$. Subfigure (e) depicts the interactions included in the ABM, which are as follows: the creation of a lost follower occurs with a probability of $1 - p_{\text{so}}$ during a head-on encounter and with a probability rate of $\lambda$ during a non-head-on encounter; in both scenarios, a minimum time $T_l$ is required to create a lost ant. The switching over happens with a probability of $p_{so}$ after time $T_l$. The returning leader turns back with a probability rate of $\rho$ after time $T_r$, and the recruitment of the lost follower occurs with a probability rate of $p_{lfr}$ as soon as a returning leader encounters a lost ant.

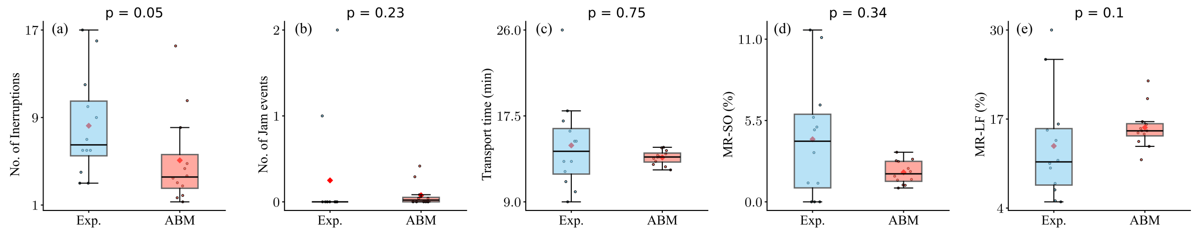


**Fig. S2.** Statistical resemblance between observations from multi-lane experiments and their respective simulations: We have depicted the data generated in simulations and observed from experiment through box-and-whisker diagrams with outliers. Subfigures $(a)$, $(b)$, $(c)$, $(d)$ and $(e)$, respectively, portray the data for number of interruptions, number of jams, total transport time, percentage of the total tandem runs involved in switching over and percentage of the total tandem runs involved in recruiting a lost ant. To examine statistical resemblance between two data sets, we have done the Mann–Whitney $U$ test and the respective $P$-values are mentioned on top of the each subfigure.

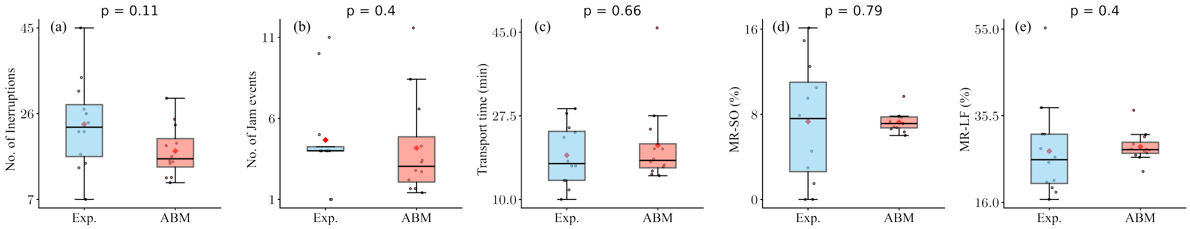


**Fig. S3.** Statistical resemblance between observations from one-lane experiments and their respective simulations: We have depicted the data generated in simulations and observed from experiment through box-and-whisker diagrams with outliers. Subfigures $(a)$, $(b)$, $(c)$, $(d)$ and $(e)$, respectively, portray the data for number of interruptions, number of jams, total transport time, percentage of the total tandem runs involved in switching over and percentage of the total tandem runs involved in recruiting a lost ant. To examine statistical resemblance between two data sets, we have done the Mann–Whitney $U$ test and the respective $P$-values are mentioned on top of the each subfigure.

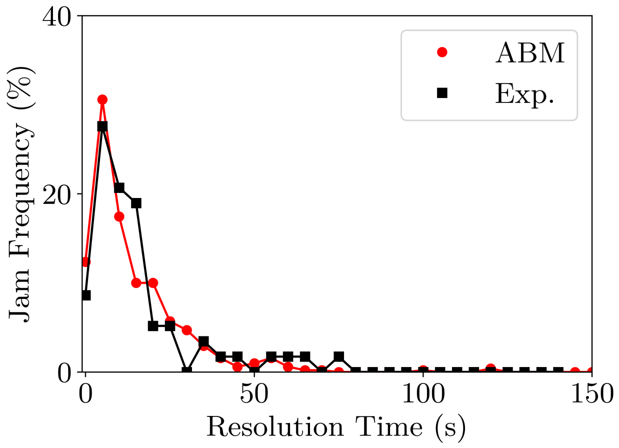


**Fig. S4.** Distribution of jam resolution time observed in experiments and generated from simulations are statistically identical (Mann-Whitney U test $P = 0.31$): The $x$ axis represents the time taken to resolve a jam in seconds and the $y$ axis represents the percentage of jams that have been resolved.

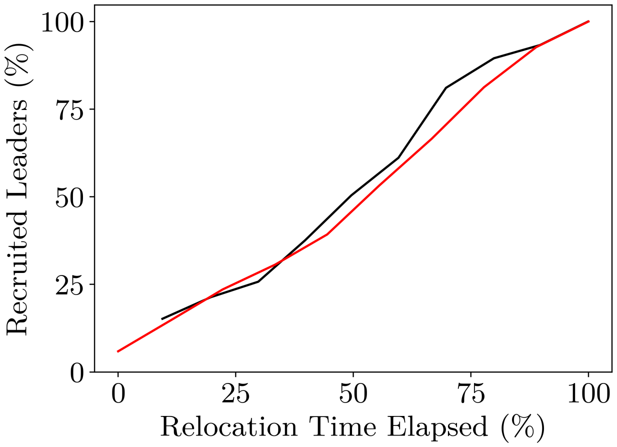


**Fig. S5.** Simulation replicates documented (1) leader recruitment dynamics: Black line and red line, respectively, correspond to leader recruitment dynamics observed in direct experiment and in simulations.

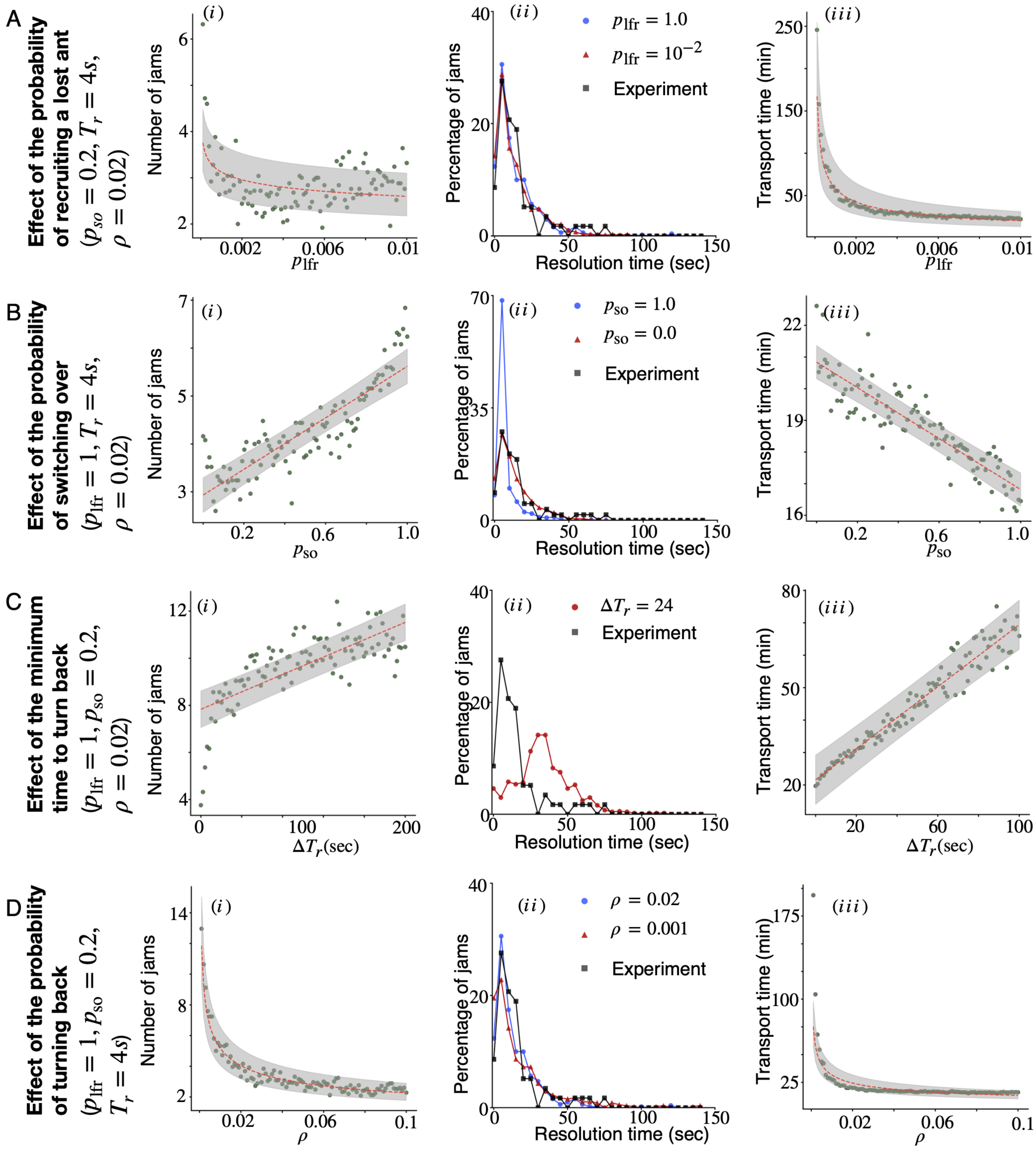


**Fig. S6.** Role of returning leaders' behavioral flexibilities in jam resolution: Each panel demonstrate how (i) the number of jams, (ii) the distribution of jam resolution time, and (iii) the transport time depend on a specific behavior of the returning leader. Panels (A), (B), (C), and (D) show the effect of the probability of recruiting a lost follower ($p_{lfr}$), the probability of switching over ($p_{so}$), the minimum time required to turn back ($T_r$), and the probability of turning back ($\rho$) after waiting for $T_r$ time , respectively. To examine the role of a specific behavior (say, switching over), we fixed all other variables ($p_{lfr}$, $\rho$ and $T_r$) as mentioned along y axis while continuously varying the parameter defining that behavior ($p_{so}$, in this case). Additionally, the gray areas in all subplots of the first and third columns represent the 95 $\%$ confidence intervals of the fitted estimate (red dashed line). These results indicate that the probability of switching over ($p_{so}$), minimum time required to turn back ($T_r$), and probability of turning back ($\rho$) are crucial factors in jam resolution, whereas the probability of recruiting a lost follower ($p_{lfr}$) plays no significant role. Furthermore, the number of jams and total transport time are influenced by all behaviors except switching over. For simulations, we have taken the size of ant colony as $87$ and averaged the data over $25$ different realizations.

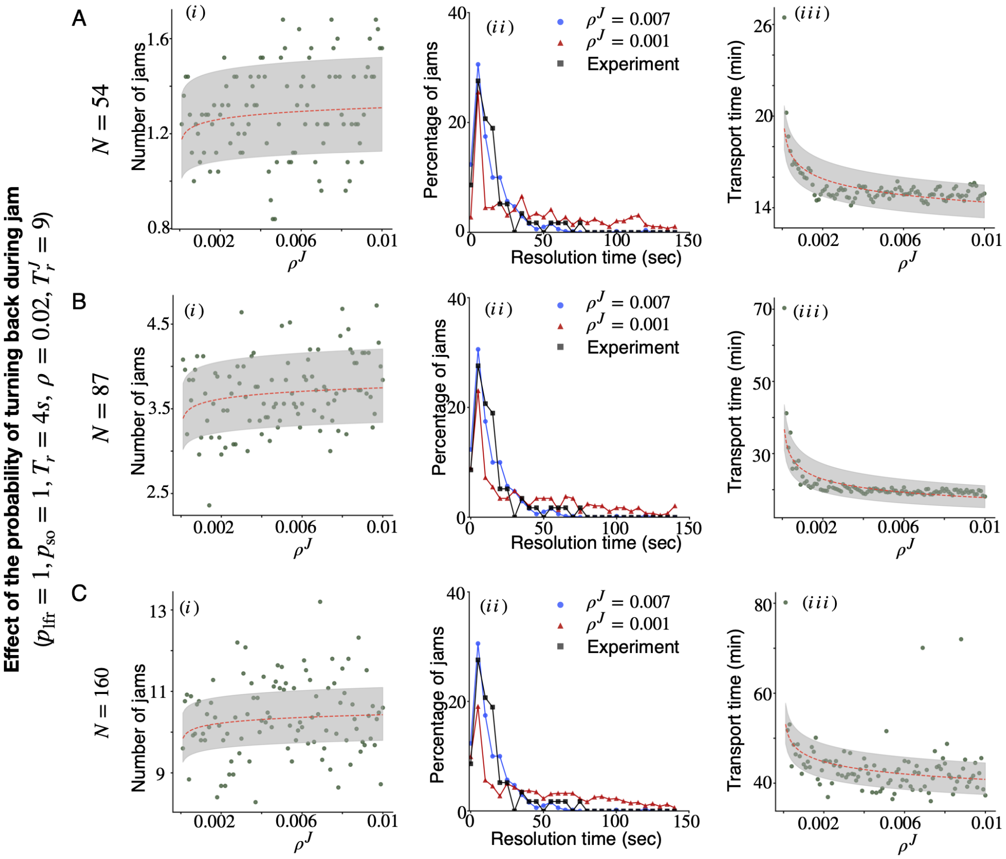


**Fig. S7.** Dependence of the relocation process on the decision of the returning leader to return towards the new nest during a jam: Subfigure (a) shows the number of jams does not depend on $\rho^J$. Subfigure (b) shows that as $\rho^J$ decreases, the fraction of jams requiring more time for their resolution increases. Lastly, subfigure (c) shows that the transport time increases as $\rho^J$ decreases. For simulations, we have taken the size of ant colonies as $54$, $87$ and $160$ (which are given in the panels A, B and C respectively) and averaged the data over $25$ different realizations.

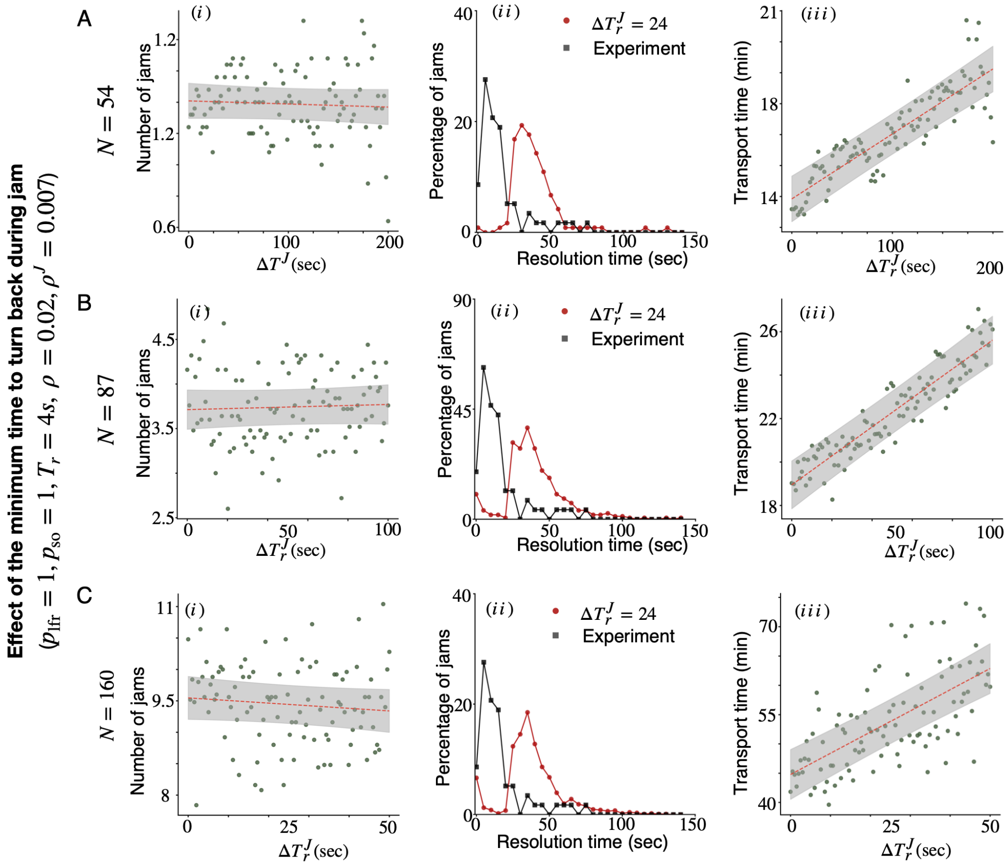


**Fig. S8.** Effect of waiting time before turning back while being stuck in a jam: Subplot (i) shows the number of jams almost independent of increment in the waiting time, $T_r^J$, by $\Delta T_r^J$. $\Delta T_r^J$, measured in seconds, is the increment in $T_r^J$ associated with the waiting times exclusively during jam scenarios before returning leader's turning back. Subplot (ii) shows that the peak of the distribution of the jam resolution time shifts towards the modified value of $T_r^J$, i.e. $9 + \Delta T_r^J = 33$. Lastly, subplot (iii) shows that the transport time increases as $T_r^J$ increases. For simulations, we have taken the size of ant colonies as $54$, $87$ and $160$ (which are given in the panels A, B and C respectively) and averaged the data over $25$ different realizations.

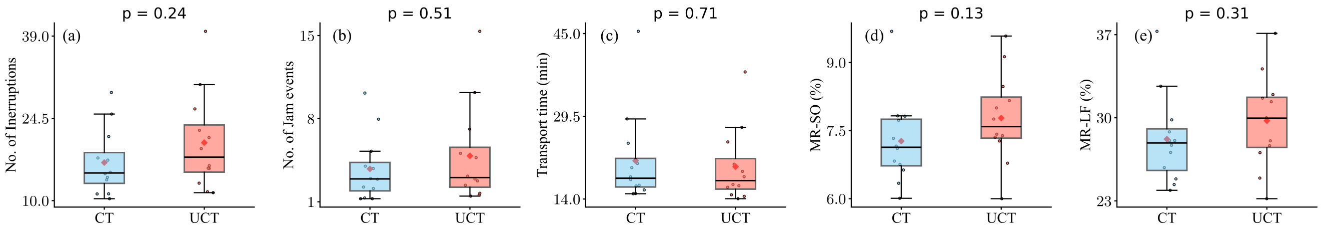


**Fig. S9.** Statistical comparison between observations from the multi-lane simulation with $coordinated\ turning$ (CT) and $uncoordinated\ turning$ (UCT) protocol: We have depicted the data generated in simulations with CT and UC protocol using box-and-whisker diagrams with outliers. Subfigures $(a)$, $(b)$, $(c)$, $(d)$ and $(e)$, respectively, portray the data for number of interruptions, number of jams, total transport time, percentage of the total tandem runs involved in switching over and percentage of the total tandem runs involved in recruiting a lost ant. To examine statistical resemblance between two data sets, we have done the Mann–Whitney $U$ test and the respective $P$-values are mentioned on top of the each subfigure.

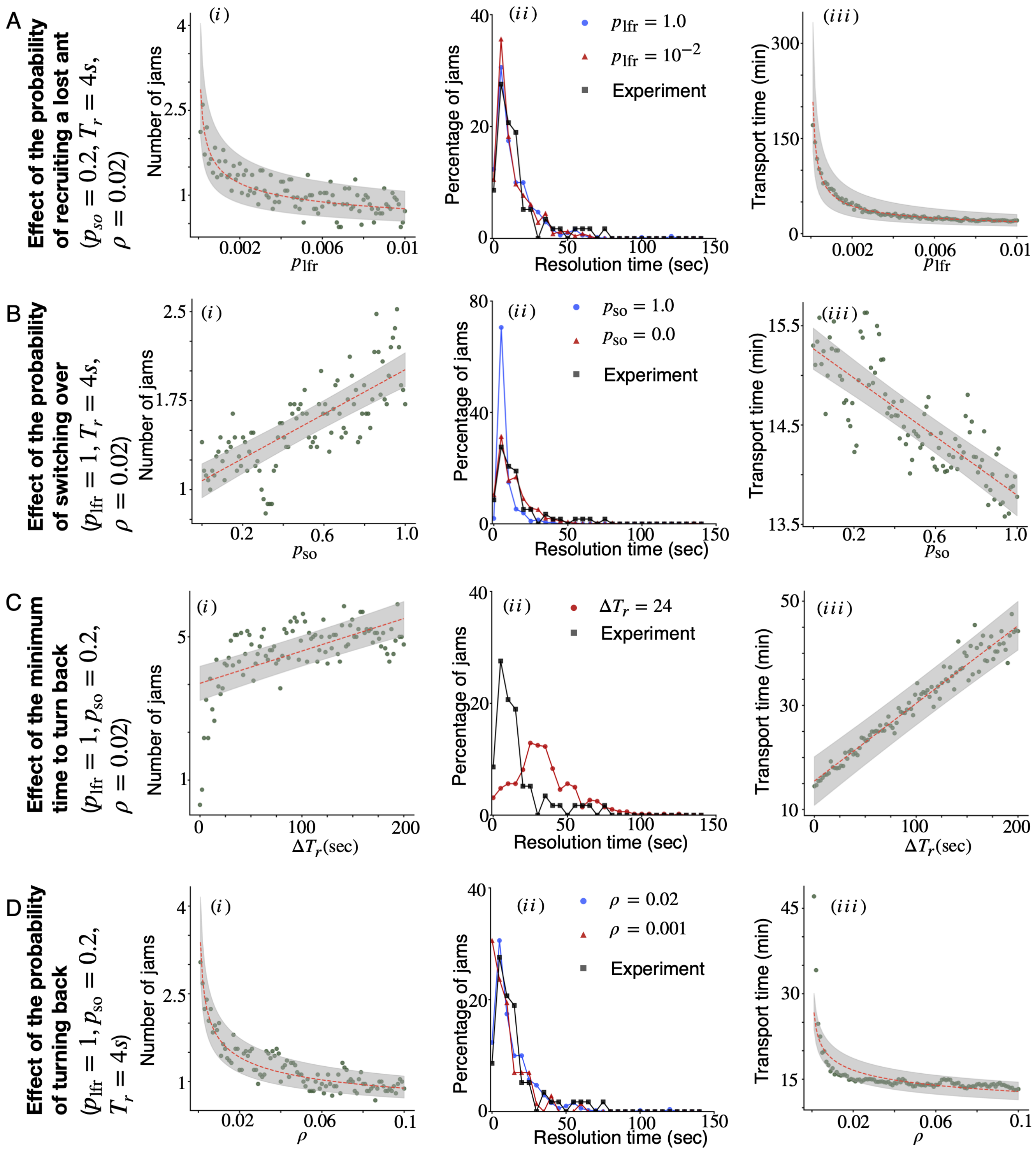


**Fig. S10.** Role of returning leaders' behavioral flexibilities in jam resolution: Each panel demonstrate how (i) the number of jams, (ii) the distribution of jam resolution time, and (iii) the transport time depend on a specific behavior of the returning leader. Panels (A), (B), (C), and (D) show the effect of the probability of recruiting a lost follower ($p_{lfr}$), the probability of switching over ($p_{so}$), the minimum time required to turn back ($T_r$), and the probability of turning back ($\rho$) after waiting for $T_r$ time , respectively. To examine the role of a specific behavior (say, switching over), we fixed all other variables ($p_{lfr}$, $\rho$ and $T_r$) as mentioned along y axis while continuously varying the parameter defining that behavior ($p_{so}$, in this case). Additionally, the gray areas in all subplots of the first and third columns represent the 95 $\%$ confidence intervals of the fitted estimate (red dashed line). These results indicate that the probability of switching over ($p_{so}$), minimum time required to turn back ($T_r$), and probability of turning back ($\rho$) are crucial factors in jam resolution, whereas the probability of recruiting a lost follower ($p_{lfr}$) plays no significant role. Furthermore, the number of jams and total transport time are influenced by all behaviors except switching over. For simulations, we have taken the size of ant colony as $54$ and averaged the data over $25$ different realizations.

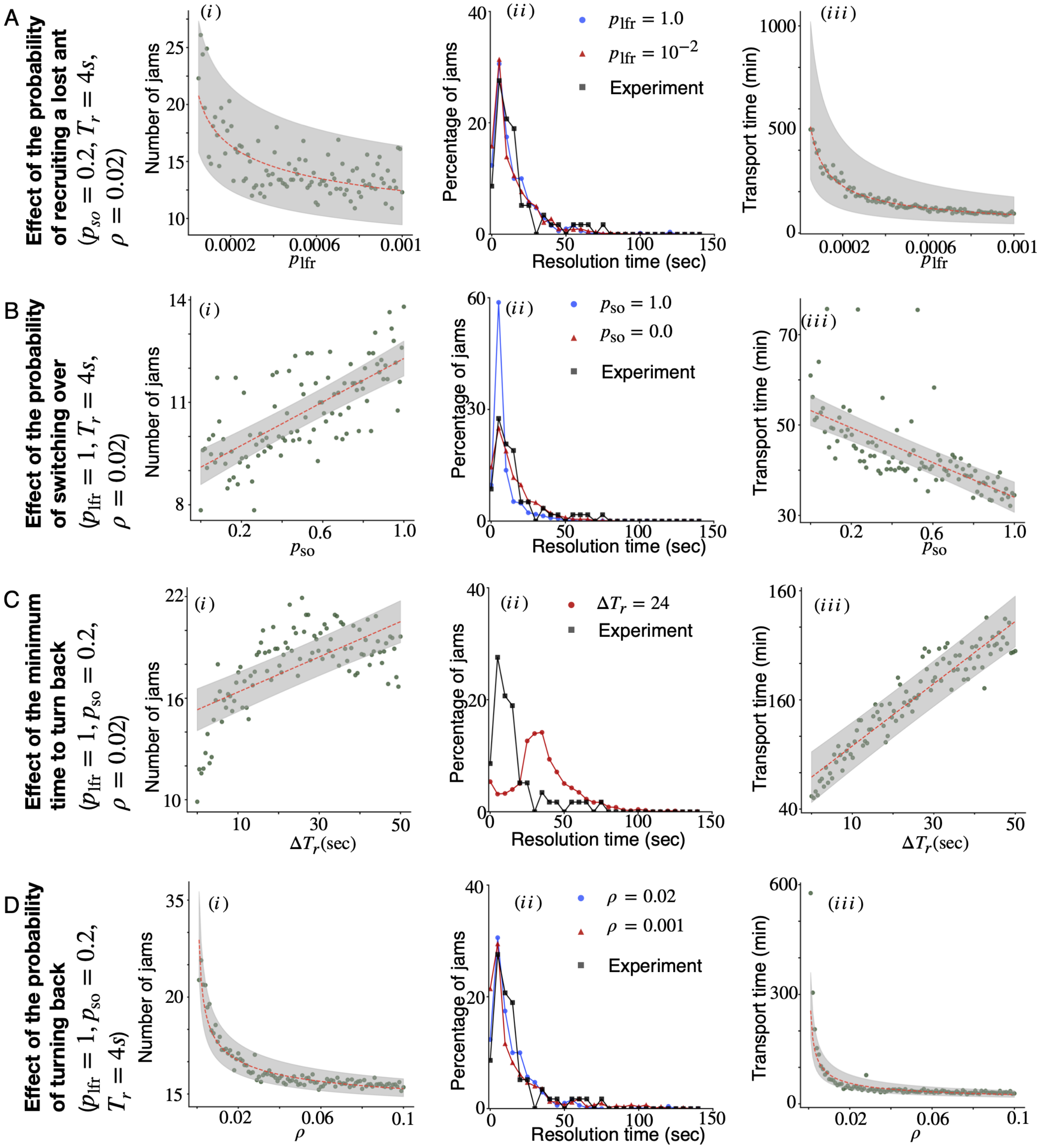


**Fig. S11.** Role of returning leaders' behavioral flexibilities in jam resolution: Each panel demonstrate how (i) the number of jams, (ii) the distribution of jam resolution time, and (iii) the transport time depend on a specific behavior of the returning leader. Panels (A), (B), (C), and (D) show the effect of the probability of recruiting a lost follower ($p_{lfr}$), the probability of switching over ($p_{so}$), the minimum time required to turn back ($T_r$), and the probability of turning back ($\rho$) after waiting for $T_r$ time , respectively. To examine the role of a specific behavior (say, switching over), we fixed all other variables ($p_{lfr}$, $\rho$ and $T_r$) as mentioned along y axis while continuously varying the parameter defining that behavior ($p_{so}$, in this case). Additionally, the gray areas in all subplots of the first and third columns represent the 95 $\%$ confidence intervals of the fitted estimate (red dashed line). These results indicate that the probability of switching over ($p_{so}$), minimum time required to turn back ($T_r$), and probability of turning back ($\rho$) are crucial factors in jam resolution, whereas the probability of recruiting a lost follower ($p_{lfr}$) plays no significant role. Furthermore, the number of jams and total transport time are influenced by all behaviors except switching over. For simulations, we have taken the size of ant colony as $160$ and averaged the data over $25$ different realizations.

#### Supplementary video 1:

**Traffic Jams and its resolution in Tandem-running *D.* indicum ants.** This video from our experiment showcases the dynamics of traffic flow in *D. indicum* ants during nest relocation along a constrained path. It displays how jams are caused by leaders returning from the old nest, who interrupt the flow towards the new nest and create traffic jams. After ants wait in the jam for multiple seconds, mass orientation towards the new nest occurs with majority of ants aligning towards the new nest to clear the jam. Events of returning leaders actively recruiting nestmates on this congested path can also be seen. All of these sections are subtitled in the top row of the video.
